# Structural insight into host plasma membrane association and assembly of HIV-1 Matrix protein

**DOI:** 10.1101/2021.02.21.432153

**Authors:** Halilibrahim Ciftci, Hiroshi Tateishi, Kotaro Koiwai, Ryoko Koga, Kensaku Anraku, Kazuaki Monde, Çağdaş Dağ, Ebru Destan, Busra Yuksel, Esra Ayan, Gunseli Yildirim, Merve Yigin, F. Betul Ertem, Alaleh Shafiei, Omur Guven, Sabri O. Besler, Raymond G. Sierra, Chun Hong Yoon, Zhen Su, Mengling Liang, Burcin Acar, Turkan Haliloglu, Masami Otsuka, Fumiaki Yumoto, Mikako Fujita, Toshiya Senda, Hasan DeMirci

## Abstract

HIV-1 continues to be a global health concern since AIDS was first recognized by the World Health Organization (WHO). It is estimated that there were 38 million people infected with HIV-1 and 1.5 million deaths in 2019 alone. A better understanding of the details of the HIV late-stage life cycle, involving Pr55^Gag^ attachment to the membrane for the further oligomerization to release virion, will provide us new avenues for potential treatment. Inositol hexakisphosphate (IP6) is an abundant endogenous cyclitol molecule and its binding was linked to the oligomerization of Pr55^Gag^ via the MA domain. However, the binding site of IP6 on MA was unknown and the structural details of this interaction were missing. Here, we present three high-resolution crystal structures of the MA domain in complex with IP6 molecules to reveal its binding mode. Additionally, extensive Differential Scanning Fluorimetry analysis combined with cryo- and ambient-temperature X-ray crystallography and computational biology identify the key residues that participate in IP6 binding. Our data provide novel insights about the multilayered HIV-1 virion assembly process that involves the interplay of IP6 with PIP2, a phosphoinositide essential for the membrane binding of Pr55^Gag^. IP6 and PIP2 have neighboring alternate binding sites within the same highly basic region (residues 18-33). This indicates that IP6 and PIP2 bindings are not mutually exclusive and may play a key role in coordinating virion particles’ membrane localization. Based on our three different IP6-MA complex crystal structures, we propose a new model that involves the IP6 coordination of the oligomerization of outer MA and inner CA domain 2D layers during assembly and budding.

## INTRODUCTION

Viruses are one of the most harmful pathogens to humanity. With their limited number of genes, they can tactfully utilize the host-cell machinery for replication via interaction between viral proteins and host-cell targets ^1^. Human immunodeficiency virus type 1 (HIV-1), which carries only nine genes, replicates in T4-helper cells ^2^. Thus, it causes acquired immunodeficiency syndrome (AIDS) by utilizing the host cell’s compartments in each step of its life cycle. One of the HIV-1 genes, *group-specific antigen* (*gag*), encodes polyprotein precursor (Pr55^Gag^) which initiates the assembly of the viral proteins and viral genomic RNA for budding of virions and contains the p6 domain that hijacks the host’s requisite proteins (Fig. 1). Virus release has i) Gag membrane binding ii) Gag-assembly and iii) the budding process, encompassing multimerization of the viral proteins and their assembly at the host cellular membrane. This process is the recruitment of key host proteins to make the viral budding and facilitates the recruitment of other essential components necessary for viral infectivities such as lipids and nucleic acids ^3, 4^. Therefore, Pr55^Gag^ is an important molecular target given what we know about its function.

**Fig. 1:**
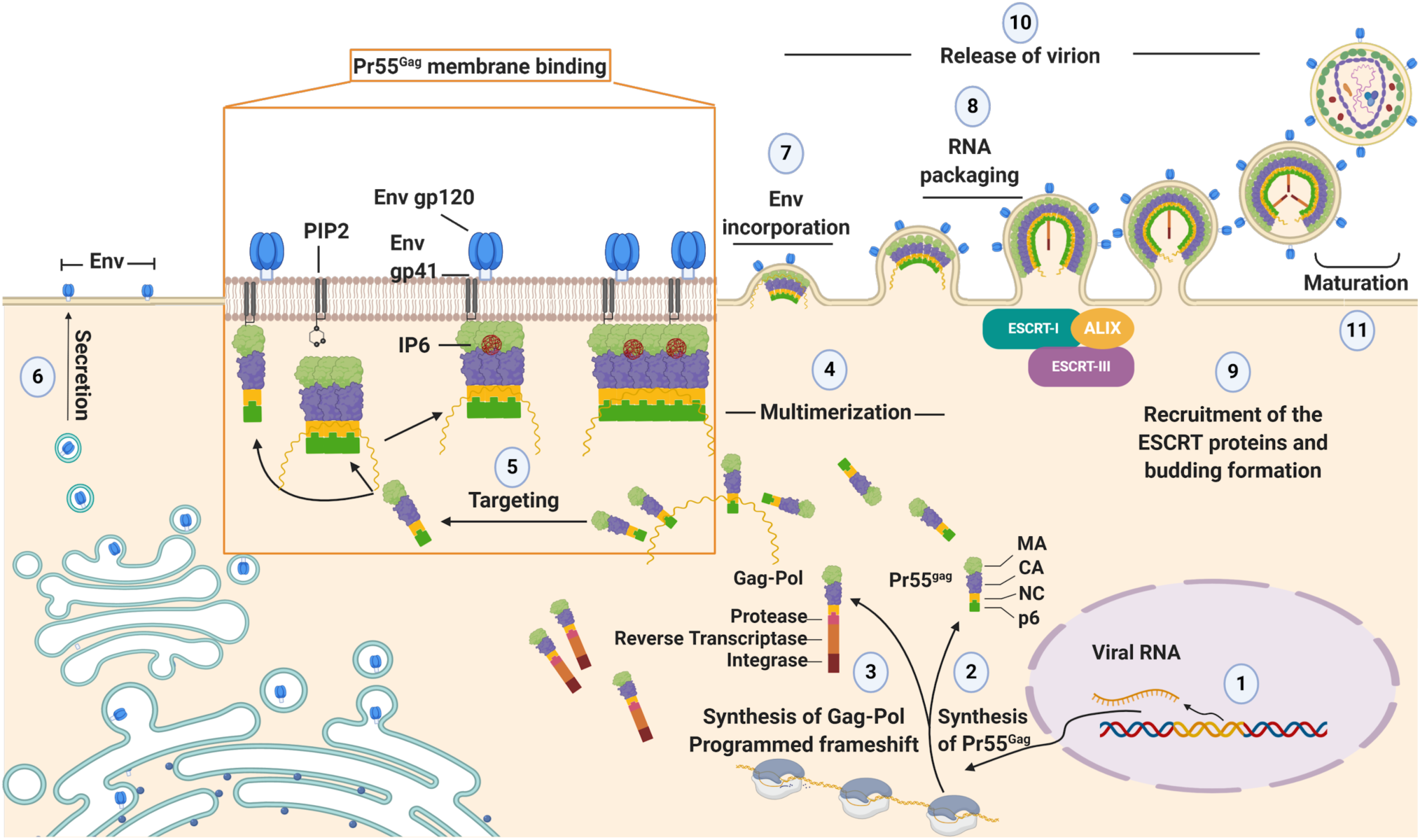
Representation of the late-stage of the HIV-1 cycle. The viral RNA is transcribed and exported from the nucleus to the cytoplasm (1). It acts as a template for the translation of Pr55^Gag^ precursor polyproteins (2). The Gag-Pol polyprotein synthesis requires a programmed frame-shifting process during translation (3). Pr55^Gag^ initiates multimerization by recruiting viral genomic RNA (4) and targets the plasma membrane for assembly through its MA domain (5). After multimerized Pr55^Gag^ polyproteins are anchored to the membrane via their amino-terminal myristate, they incorporate with Env (7), which arrives at the plasma membrane via a secretory pathway from the Golgi and RER (6). During the RNA packaging (8) and budding process, ESCRT-I and ESCRT-III are recruited to perform a membrane scission (9) which is characterized by the release of the virion particle (10). Finally, maturation (11) is catalyzed by proteolytic cleavage of the viral protease for canonical capsid core formation. **ALIX**: (ESCRT-associated factor) ALG2-interacting protein X, **CA**: capsid domain, **Env**: viral envelope glycoprotein, **ESCRT-I**: endosomal sorting complex required for transport I, **MA**: matrix domain, **NC**: nucleocapsid domain.

In conjunction with the beginning of the HIV-1 replication cycle, Pr55^Gag^ is synthesized from the viral genomic RNA transcript inside the host cell. Pr55^Gag^ consists of four domains: the matrix domain (MA), capsid domain (CA), nucleocapsid domain (NC) and p6 domain respectively. In addition, Pr55^Gag^ has two small spacer peptides SP1 between CA and NC, and SP2 between NC and p6 which influence virus release efficiency and infectivity (Supplementary Fig. 1). Pr55*^Gag^* is directed to the plasma membrane via its MA domain and is incorporated into the viral envelope (Env) glycoproteins to create virion particles ^5, 6^. Moreover, MA is crucial for the proper conformation of HIV-1 Gag protein. Approximation of MA’s N-terminal domain with the C-terminal domain of NC provides a horseshoe-like conformation of Gag protein ^7^. Maturation of the virions is the final step of the late stage of infection and is initiated by cleavage, that is, separation of Gag domains via the action of a specific viral protease ^3, 4^.

The MA domain of Pr55*^Gag^* contains five α-helices: four of which form an N-terminal globular head while the fifth α-helix forms a C-terminal helix that is connected to the CA domain ^8^. The MA domain has two signals for the Gag integration into the host plasma membrane. One of these is the “highly basic region” (HBR), which includes the residues (18-33). As the other signal, the assembly of Pr55^Gag^ at the plasma membrane requires myristoylation of the MA domain at its N-terminus ^9^. Moreover, 1-D-*myo*-phosphatidylinositol 4,5-bisphosphate [PI(4,5)P2] (PIP2), which is a native host membrane phospholipid, is recognized via the MA domain of Pr55^Gag^ during this responsible step for the localization of membrane proteins ^10^. The HBR within the N-terminus of MA has an essential role, leading hotspot activity by providing necessary interaction between MA and PIP2 ^11, 12^. In addition, MA has different observed oligomeric states: trimer, hexamer and higher-order oligomers on the host plasma membrane ^13^. As any structural ablation of the higher-order state will promote the viral release inhibition, it makes the amino acid sequence in the N-terminus critical ^14^.

Detailed high-resolution structural studies of MA-inositol complexes are still not available and this impedes the further understanding of the assembly of Pr55^Gag^ at the plasma membrane. Unlike previous studies ^15–17^, high-resolution studies in this direction will offer invaluable opportunities for molecular designs of lead high-efficiency compounds that may inhibit the completion of the HIV cycle by blocking the Pr55Gag-PIP2 interaction ^18^. Studies such as the solution NMR structure of MA-PIP2 complex with shorter lipophilic chains revealed that a basic pocket of MA domain accommodates acidic inositol phosphate moieties by electrostatic interactions ^19^. Previous studies suggested that the other inositol molecules such as inositol hexakisphosphate (phytic acid, IP6) in addition to PIP2 are essential for the canonical HIV-1 particle assembly process ^20^. The IP6 is an abundant endogenous molecule in organisms, especially in plants and has critical roles in biological activities such as IP signaling cascade. Although it was thought previously to be present only in plants, it has been discovered that the presence of IP6 is also very common within the animal kingdom ^21^. IP6 as an abundant host metabolite has been hijacked for the benefits of HIV-1 through interaction with amino acid residues within the SP1 region of Pr55^Gag 7, 22^. Furthermore, IP6 is involved in the assembly of Pr55^Gag^ by binding to both MA and NC domains which are responsible for the recruitment of the viral genome into virions and facilitate the assembly process and encapsidation ^23^. The binding of IP6 to the CA domain increases the HIV-1 capsid stability and RNA accumulation, leading to viral production. Therefore, the endogenous concentration of IP6 is critical for HIV-1 assembly ^22^. In the presence of IP6, HIV-1 Gag performs conformational transition into virus-like particles (VLPs). The inositol phosphates (IPs) provide to correct the radius of curvature during the assembly process of Pr55^Gag^, leading to the alteration of VLPs for getting mature conformation ^7, 23^. Moreover, MA protein is a potential prime therapeutic target as it involves the assembly of the viral particles ^24^. While IP6 is a molecule that can play a role in the virus assembly by binding with the MA domain ^25^ the interaction between IP6 and the MA domain of Pr55^Gag^ has not been fully elucidated for HIV-1 progression.

It is essential to determine the high-resolution structures of the MA-IP6 complexes to obtain molecular insight into the dynamic interaction between MA and IP6 at the molecular level. Here we employed both synchrotron cryo X-ray crystallography and ambient-temperature Serial Femtosecond X-ray crystallography (SFX) at an X-ray Free Electron Laser (XFEL) to obtain MA-IP6 co-crystal structures. We report three crystal forms and resulting X-ray structures of MA in complex with IP6 molecule (MA_IP6). One of our crystal structures also reveals a novel form of a hexameric state of MA in the presence of IP6. Our data demonstrate that the binding site of MA-IP6 is different from the PIP2-binding site and through this interaction, IP6 shows significant importance in the oligomerization of MA. Our crystallographic data is also supported by extensive net Transfer Entropy (nTE) analysis of monomeric, trimeric, hexameric and dodecamer structures of MA protein. Accordingly, the IP6 binding site directly affects trimerization, PIP2 binding, envelope interaction, and myristoylation sites in a monomeric state. Moreover, IP6, PIP2, myristoylation, and C-terminal of the domain interact with trimerization sites mutually in both trimeric and hexameric states. In conclusion, our data suggest that IP6 binding of MA regulates Gag protein localization and higher-order oligomerization on the host membrane even in the monomeric or hexameric state of MA. These findings may provide a better understanding of the HIV-1 replication mechanism besides a novel insight into the interaction between MA and IP6.

## MATERIALS AND METHODS

### Reagents

IP6 was purchased by Merck (Merck, Darmstadt, Germany). All the chemicals used for crystallographic studies were ultrapure grade and purchased from Sigma-Aldrich, USA.

### Cloning and overexpression of the HIV-1 Gag MA domain

A protocol for bacterial expression and purification of the MA domain is previously described ^26^. A gene encoding MA fusion with Tobacco Etch Virus (TEV) protease cleavage site at the N-terminus site was subcloned between *Kpn*I (5’-terminus) and *Hind*III (3’-terminus) restriction enzyme sites in the pRSF-1b vector (Merck, Darmstadt, Germany) to express His-TEV protease site-MA protein (pRSF-1b_MA) (10His-tagged MA).

### Protein expression and purification

10His-tagged MA gene was expressed in *Escherichia coli* (*E. coli*) strain BL21 (DE3) (Merck, Darmstadt, Germany). The 10His-tagged MA protein was initially purified with nickel-NTA affinity column chromatography followed by TEV cleavage, denaturing and refolding, and size exclusion column chromatography. *E. coli* BL21 (DE3) transformed with pRSF-1b_MA was cultured in LB broth supplemented by 50 mg/mL kanamycin at 37 °C. When the culture reached OD_600_ of approximately 0.6, target protein expression was induced by 0.1 mM Isopropyl β-D-1-thiogalactopyranoside (IPTG) at 16 °C overnight. Harvested cells were resuspended in a lysis buffer containing 20 mM Tris-HCl (pH 8.0), 500 mM NaCl, 30 mM Imidazole, 5 mM β-mercaptoethanol and lysed by sonication (ULTRA5 HOMOGENIZER VP-305, TAITEC, Output 7, Duty 50%, 2 min). The lysate was separated into pellet and supernatant by centrifugation at 15,000 *g* for 30 min at 4°C, and the supernatant was applied to a nickel-NTA affinity column containing a total of 4 ml bed volume of nickel-NTA superflow resin (Qiagen, Venlo, Netherlands). After washing with a washing buffer containing 20 mM Tris-HCl, pH 8.0, 1 M NaCl, 1 M (NH_4_)_2_SO_4_, 30 mM Imidazole, 5 mM β-mercaptoethanol, the target protein was eluted with 15 ml of an elution buffer that contains 20 mM Tris-HCl (pH 8.0), 500 mM NaCl, 1 M (NH_4_)_2_SO_4_, 5 mM β-mercaptoethanol, 300 mM Imidazole. The eluent was concentrated to 2 ml, and 10His tag was removed by 0.1 mg/ml 6His-tagged TEV protease while dialyzed overnight in dialysis buffer containing 20 mM Tris-HCl (pH 8.0), 500 mM NaCl, 5% glycerol, 1 mM DTT. The mixture was diluted to be 50 ml, and His-tagged TEV protease, cleaved 10His tag and uncleaved 10His-tagged MA were separated by nickel-NTA affinity chromatography, and the cleaved un-tagged product was collected at the flow-through fraction of the affinity chromatography. The 2 ml of fraction was mixed with 10 ml of denaturation buffer containing 20 mM Tris-HCl (pH 8.0), 1 M NaCl, 1 M (NH_4_)_2_SO_4_, 1 mM DTT, 6 M urea and the protein was denatured overnight during dialysis in the denaturing buffer. After concentrating to a final volume of 1 ml, polypeptides were purified with size exclusion chromatography (SEC) column Superdex 200 10/300 increase (GE Healthcare, Little Chalfont, United Kingdom) which equilibrated with the SEC buffer containing 20 mM Tris-HCl (pH 8.0), 150 mM NaCl, 1 mM DTT using Akta FPLC system (GE Healthcare) at 0.2 mL/min. Peak fractions containing refolded MA were collected and concentrated to a final concentration of 10 mg/ml for crystallization.

### Crystallization for cryo synchrotron studies

200 mM Phosphatidylinositol-6-phosphate (IP6) was mixed with 10 mg/ml MA protein solution. Protein Crystallization System (PXS) was used for the initial crystallization screening of the MA-IP6 complex ^26^. In total, 384 conditions were examined using Crystal Screen 1 & 2 (Hampton Research), Index (Hampton Research), PEG-Ion (Hampton Research), and PEG-Ion2 (Hampton Research), and Wizard I & II (Molecular Dimensions). First crystallization condition contained 25% (w/v) PEG 3350 and 100 mM MES (pH 6.0) (MA_IP6_1), and the second one contained 5% (w/v) PEG 4000, 20% (v/v) 2-propanol and 100 mM MES (pH 6.5).

### Cryogenic data collection and processing

Crystals were harvested with the proper size of MicroLoops (MiTeGen, New York, United States of America), flash frozen by plunging into liquid nitrogen and transferred in Uni-pucks for automated data collection. Diffraction data were collected at the beamline BL-1A in the Photon Factory. The diffraction data sets were automatically processed and scaled using *XDS* ^27^, *POINTLESS* ^28^, and *AIMLESS* ^29^. Crystallographic statistics are summarized in Supplementary Table 1.

### Crystal structure determination and refinement of cryogenic synchrotron structures

Phases for all three MA_IP6 structures data were determined by the molecular replacement method using the crystal structure of MA (PDB ID: 1HIW) as a search model with *MOLREP* ^30^. Molecular models were initially refined with *REFMAC5* ^31^ and further refined by *PHENIX.refine* ^32^. Molecular models were manually built by *COOT* ^33^. m*F*o - D*F*c omit-maps for ligands were calculated using *PHENIX* with a simulated annealing protocol. Interaction between MA and IP6 was analyzed by *PISA* ^34^. All molecular graphics in this manuscript were prepared by the *PyMOL* Molecular Graphics System, Version 2.3.0 Schrödinger, LLC.

### Batch crystallization of the HIV-1 Gag MA domain for SFX

Purified HIV-1 Gag MA proteins were used in co-crystallization with IP6 at room temperature by the hanging-drop method using a crystallization buffer containing 20% (w/v) polyethylene glycol 3350 (PEG 3350) as a precipitant and 100 mM MES-NaOH (pH 6.5) ^35^. Microcrystals were harvested in the same mother-liquor composition, pooled to a total volume of 3 ml and filtered through a 40-micron Millipore mesh filter. The concentration of crystal was around 10^10^-10^11^ per milliliter viewed by light microscopy.

### X-ray Free Electron Laser data collection parameters

An average of 2.64 mJ was delivered in each 40-fs pulse containing approximately 10^12^ photons with 9.51 keV photon energy with 1×1 mm^2^ focus of X-rays. Single-pulse diffraction patterns from HIV-1 MA-IP6 microcrystals were recorded at 120 Hz on a CSPAD ^36^ detector positioned at a distance of 217 mm from the interaction region.

### Sample delivery of MA-IP6 microcrystals into an XFEL and data collection

A crystalline slurry of MA-IP6 microcrystals kept at ambient-temperature flowing at 2 µl/min was injected into the interaction region inside the front vacuum chamber at the LCLS CXI instrument using the coMESH injector ^37^. Large size crystals were present in the MA-IP6 sample slurry, prior to the experiment. The coMESH injector required filtered samples before injection through a 100-micron inner diameter capillary size to prevent clogging.

### Hit finding and indexing

SFX diffraction data collected from rod-shaped crystals at LCLS were processed using *Psocake* 38, ^39^, yielding a complete dataset. A diffraction pattern was deemed a hit if at least 15 peaks were found. A total of 126,434 diffraction patterns for MA-IP6 microcrystals were recorded as crystal hits. *CrystFEL*’s *indeximajig* program ^40, 41^ was used to index the crystal hits (Supplementary Fig.2). Two rounds of indexing were performed on the dataset. Initial indexing results indicated that the space group was most likely triclinic P1 with a = 96.73 Å, b = 96.97 Å, c = 91.76 Å and α = 90.05°, β = 90°, γ = 120°. Given the target unit cell, the indexing results were accepted if the unit cell lengths and angles were within 5% and 1.5°, respectively. The final iteration yielded 56,861 (45%) indexed patterns for the MA-IP6 dataset.

### Differential Scanning Fluorimetry assay

A protocol for Differential Scanning Fluorimetry (DSF) was employed according to the previous procedures ^42, 43^. SYPRO^®^ Orange (Thermo Fisher Scientific, Massachusetts, United States of America) solution (× 5000) was diluted by 50 times with a buffer containing 50 mM MOPS-NaOH (pH 7.0), 50 mM NaCl (100 × SYPRO^®^ Orange solution). A buffer solution containing purified His10-tagged MA protein was exchanged to the 50 mM MOPS-NaOH, 50 mM NaCl buffer (pH 7.0) with ultrafiltration using Amicon^®^ Ultra-15 Centrifugal Filter Unit (10 kDa MWCO) (Merck, Darmstadt, Germany). With the prepared 100 × SYPRO^®^ Orange solution and the protein solution, 50 μl of a reaction mixture were prepared to be 5.4 µM MA-His10 protein, 5 × SYPRO^®^ Orange in presence of or in absence of the indicated concentration of compounds to evaluate the contribution of IP6 to the thermostability of MA protein, or 32.4 µM IP6 for MA point-mutant analysis. The temperature of the reaction system was increased 0.5°C per 30 sec from 20 to 95°C stepwise, and fluorescence intensity at each temperature was measured by Single-Color Real-Time PCR Detection System, MyiQ (Bio-Rad, California, United States of America), and analyzed with iQ5 (Bio-Rad).

### Gaussian Network Model (GNM)-Based Transfer Entropy Calculations

The Gaussian Network Model (GNM) is the most minimalist isotropic elastic network model of Cα atoms with harmonic interactions for the dynamics of proteins and their complexes ^44, 45^. GNM-based TE calculations ^46, 47^ reveal causal interrelations of residues in a given structural topology by considering a certain time delay τ between residue fluctuations *ΔR*_i_ (t) of residue i at time t and *ΔR*_j_ (t+**τ**) of residue j at time t+**τ**. Using these fluctuations, TE(i,j) (**τ**) provides an estimate for the direction of information flow from residue i to residue j in time delay **τ**; i.e. TE(i,j) (**τ**) describes how much the present movement of residue i decreases the amount of uncertainty for the future movement of residue j within time interval **τ**. If TE(i,j)(**τ**) > TE(j,i)(**τ**), the dynamics of residue i affects the dynamics of residue j, indicating a causal directional interrelation from residue i to j in time delay **τ**.

The transfer entropy TE(i,j)(τ) from each residue pairs of i at time t and j at time t+**τ** was formulated^46^ as:

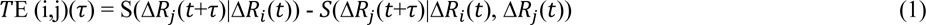

where the conditional entropies are:

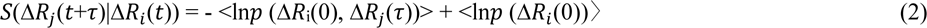

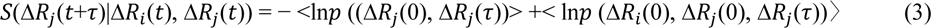

Each term in Equations 2 and 3 could be obtained using GNM ^46^, which mainly bases the calculations on the topology of the structure by computing the connectivity/ Kirchoff matrix from cartesian coordinates. Time delay **τ** between fluctuations Δ*R*_*i*_ and Δ*R*_*j*_ is used as the four-fold of the defined time variable which maximizes mean transfer entropies over all residue pairs i and j.

GNM decomposes the motions into a spectrum of dynamic modes, from global/slow (low frequency) modes to local/fast (high frequency) modes. TE values in Equation 1 could be calculated using all or a subgroup of dynamic modes ^47^. As global modes of motion are highly relevant for functional dynamics ^48^, here we considered the slowest end of the dynamic mode spectrum with and without the slowest mode; average ten slowest and average two to ten slowest dynamic modes.

To emphasize the dynamically affecting and affected residues of each pair i and j, net TE (nTE) values is defined as

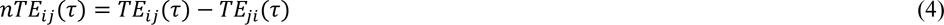

nTE is displayed with color (from blue to red as transfer entropies ascend) for each residue pair i and j on transfer entropy heat maps (Supplementary Fig. 6-10). For representative residues (highlighted in grey), their causal interrelation with the rest of residues is color-coded from highest positive (red) to lowest negative (blue) nTE values on all used topologies, given next to the nTE heat maps.

To identify affecting residues as entropy sources with the capacity to affect the others at the most, i.e. transfer information to the others, among which the maximally affected residues are defined as entropy sinks, the cumulative net transfer entropy (-’”() values of residue i over all j residues were also calculated as

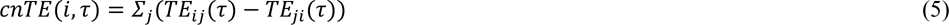

*cnTE* values of residue pairs are given as a line plot on the left of transfer entropy heatmaps (Supplementary Fig. 6-10).

Using the first ten global modes reveals the main network of bidirectional causal interrelations among IP6 and PIP2 binding sites, envelope protein and myristoyl interaction sites, trimeric interfaces and further oligomerization sites of MA domains. The exclusion of the slowest mode discloses the subtler causal interactions. Please note that IP6 molecules were not considered explicitly in the calculations, which thus mostly reflect the intrinsic behavior of monomer and oligomers.

## RESULTS

### Crystal structures of MA-IP6 complexes

We employed X-ray crystallography to reveal the molecular mechanism of interaction between the MA domain and IP6 molecules. MA protein and IP6, which is the most abundant inositol *in vivo*, were co-crystallized. From three crystal forms, different crystal structures of MA complexed with IP6 (MA_IP6) were determined. Cryogenic MA_IP6_R32 structure is at 2.40 Å, Cryogenic MA_IP6_C2 extends to 2.72 Å, and ambient-temperature MA_IP6_SFX structure is determined at 3.5 Å resolution, respectively (Fig. 2 & 3, Supplementary Fig. 3). Our overall MA_IP6 complex structures are consistent with the previous NMR and crystal structures (PDB IDs: 2HMX and 1HIW, respectively) with the MA domain composed of 5 alpha-helices (Supplementary Table 2 & 3). Compared to the previous structures, a globular domain composed of the 1st to 4th helices did not show significant conformational change, whereas the C-terminal 5th helix did and will be further discussed later. Trimeric assembly of MA showed no significant overall structural differences between the MA_IP6 complexes and MA Apo form (PDB ID: 1HIW). However, a superposition of chain A of MA_IP6_C2 and 1HIW form showed side-chain conformational changes within the HBR (18-33) of the N-terminus specifically residues Lys18, Arg20, Arg22, Lys26, Lys27, Gln28, Lys30, Leu31, Lys32, and His33 (Fig. 4). These conformational changes are involved in the interaction between MA and IP6 (Arg20, Gln28, Lys18) in our monomeric structure. Based on the density map of MA_IP6_C2, the *B*-factors of each IP6 molecule are notably high compared to MA (Fig. 3 & Supplementary Table 4). This suggests that the binding modes of IP6 molecules are not well-defined and that the affinities of MA with each IP6 varies. In our MA_IP6_R32 crystal form, the density map derived from IP6 was also observed at the same binding sites (Fig. 2). The observed differences of individual IP6 molecule interactions in each MA structure will be further explained below.

**Fig. 2:**
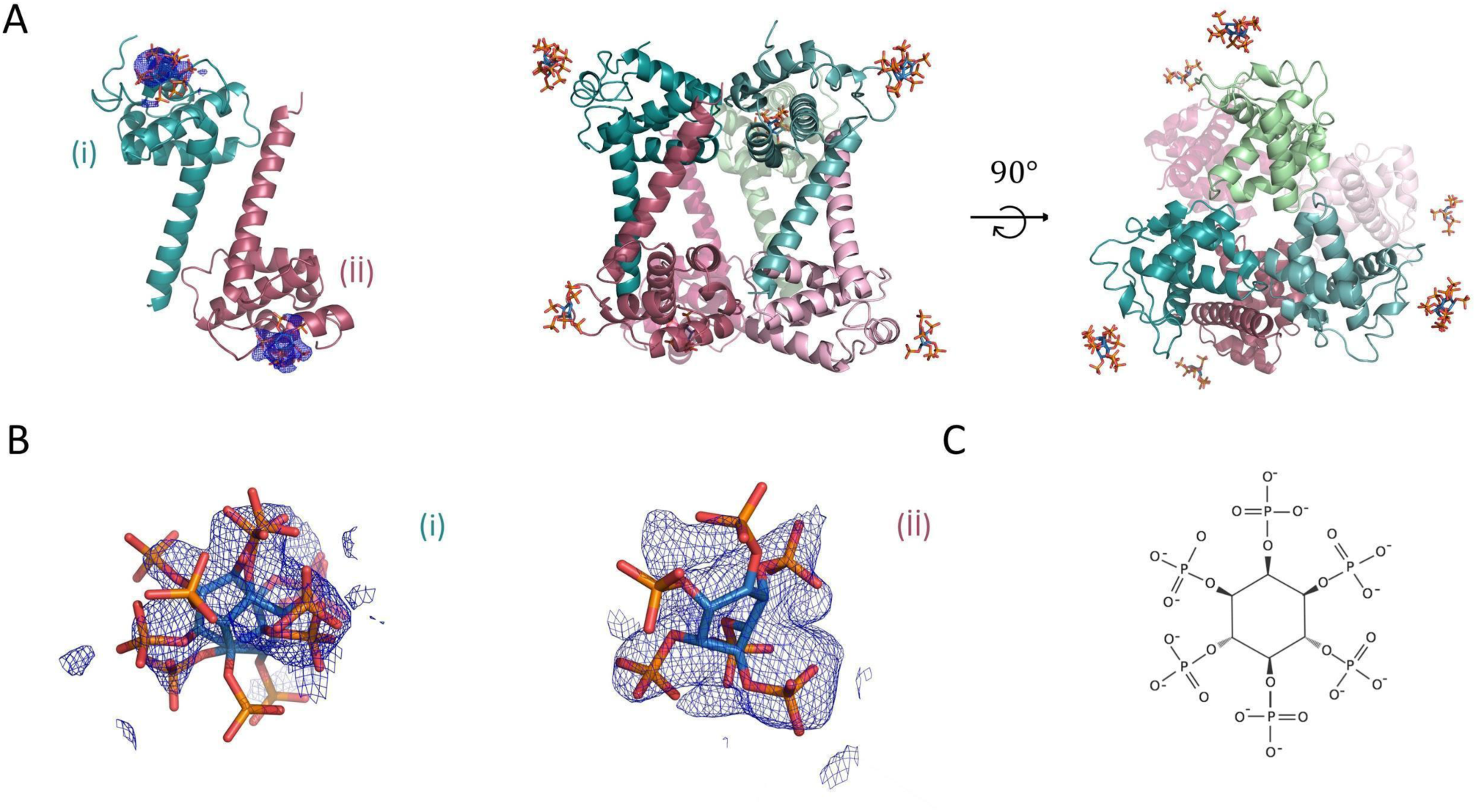
The overall structure of MA_IP6_R32 with symmetric units. **a** Chains A and B of MA_IP6_R32 structure in the asymmetric unit cell are colored in deep-teal and raspberry, respectively. Chain A and B symmetry mates are generated in *PyMOL* and are colored in pale-green and light-pink for the representation of symmetric units, respectively. Carbon, oxygen and phosphorus atoms of IP6 molecules are colored by sky-blue, red and orange, respectively. **b** 2*F*o-*F*c simulated annealing-omit map is colored in blue mesh and shown at 2.5σ level within 5.0 Å from IP6. **c** Chemical structure representation of the *myo*-IP6 molecule.

**Fig. 3:**
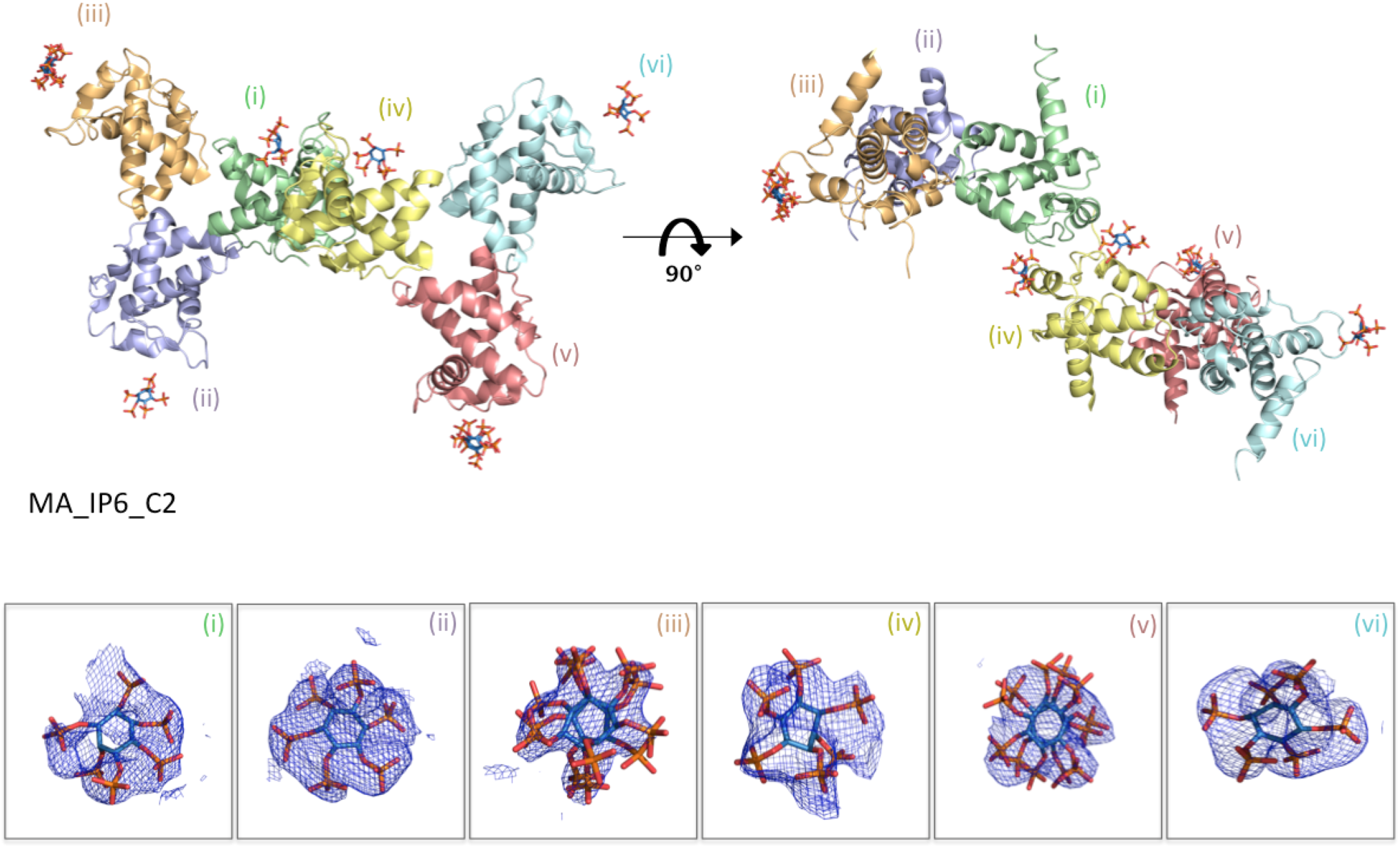
The overall structure of MA_IP6_C2 in an asymmetric unit. MA proteins in the asymmetric unit are shown by cartoon representation. Each chain is indicated by a different color and labeled with the respective color (pale-green: chain i, light blue: chain ii, light-orange: chain iii, pale-yellow: chain iv, salmon: chain v, pale-cyan: chain vi). Carbon, oxygen and phosphorus atoms of IP6 are colored by sky-blue, red and orange, respectively. A 2*F*o-*F*c simulated annealing-omit map colored in blue mesh at 2.5σ level within 5.0 Å from IP6 is shown. For better view density maps derived from IP6 molecules are enlarged and shown in the insets.

**Fig. 4:**
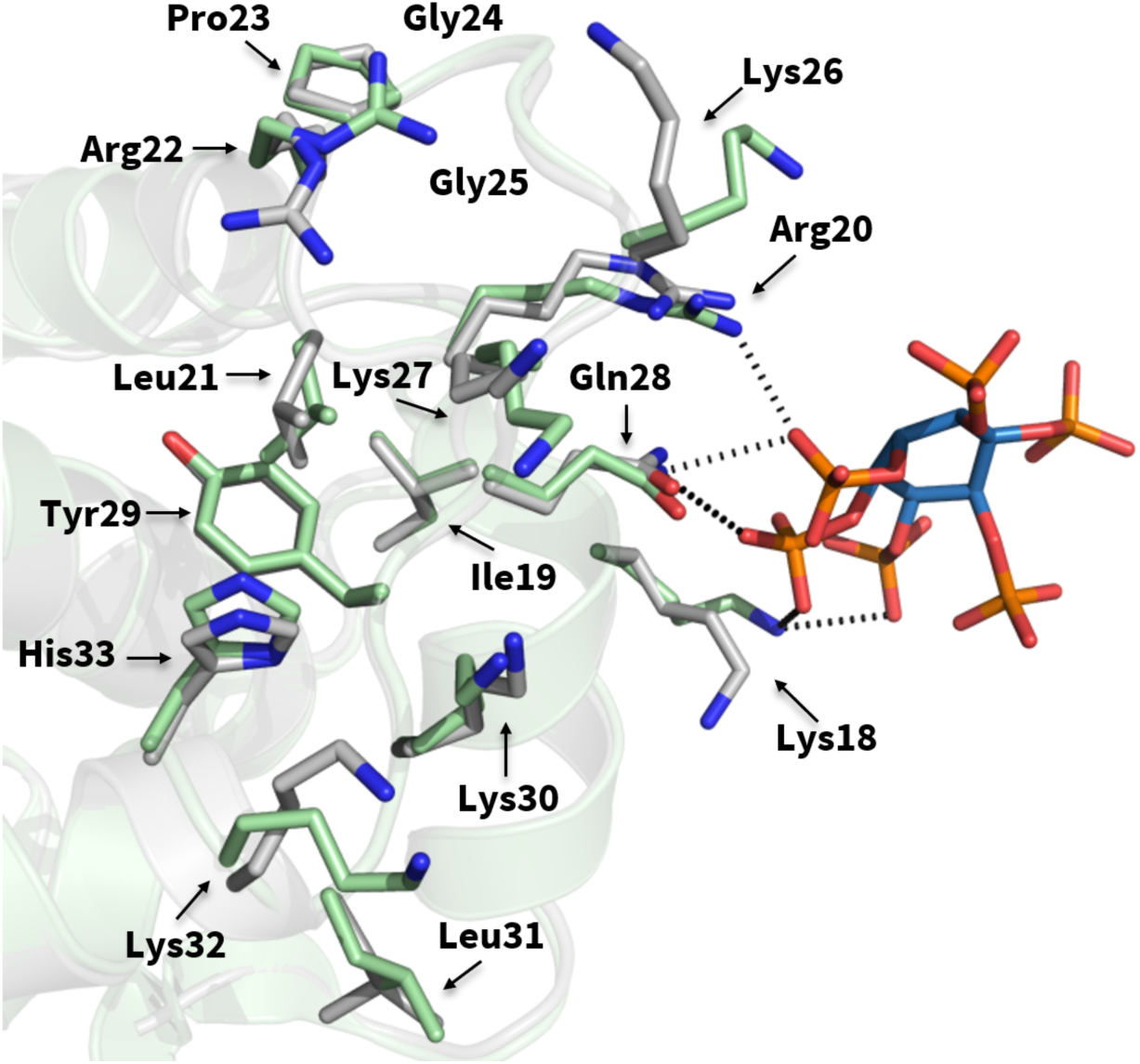
Superposition of MA domain structure (PDB ID: 1HIW) and our MA_IP6_C2 structure. The chain A of our structure in green is superposed with chain A of MA domain (PDB ID: 1HIW) in gray, RMSD 0.279 Å. The conformational changes on the residues in the HBR (18-33) are indicated and hydrogen bonds within the MA_IP6_C2 complex are shown with dotted lines.

### IP6-binding sites of MA

In the MA_IP6_C2 structure, a single chain of oligomeric MA interacts with three IP6 molecules. Three IP6 molecules were observed on chain A (Fig. 5A-E); one is on the asymmetric molecule of chain A (Fig. 5E, IP6_1), another is mainly in contact with chain D and visualized on chain A (Fig. 5D, IP6_2), and the last one is on the asymmetric molecule of chain A (Fig. 5C, IP6_3). Lys18, Arg20 and Glu28 formed hydrogen bonds with IP6_1. Lys27, Lys30, Lys32 and His33 are located close to IP6_2 and two hydrogen bonds were observed between Lys27, Lys30 and IP6_2. Ala3, Arg4 and Arg39 are near IP6_3 and Ala3, Arg4 interacted with this IP6 molecule. Additionally, when the MA_IP6_R32 structure was superposed with the MA_IP6_C2 structure, the binding position of IP6 molecules in each chain was detected in the same binding surface for both crystal forms (Supplementary Fig. 4). The interactions between IP6 and two crystal forms were analyzed to investigate the different non-covalent interactions, but the same interacted residues were observed for both structures (Supplementary Fig. 5). To evaluate and dissect the contribution of these interactions, we employed Differential Scanning Fluorimetry (DSF).

**Fig. 5:**
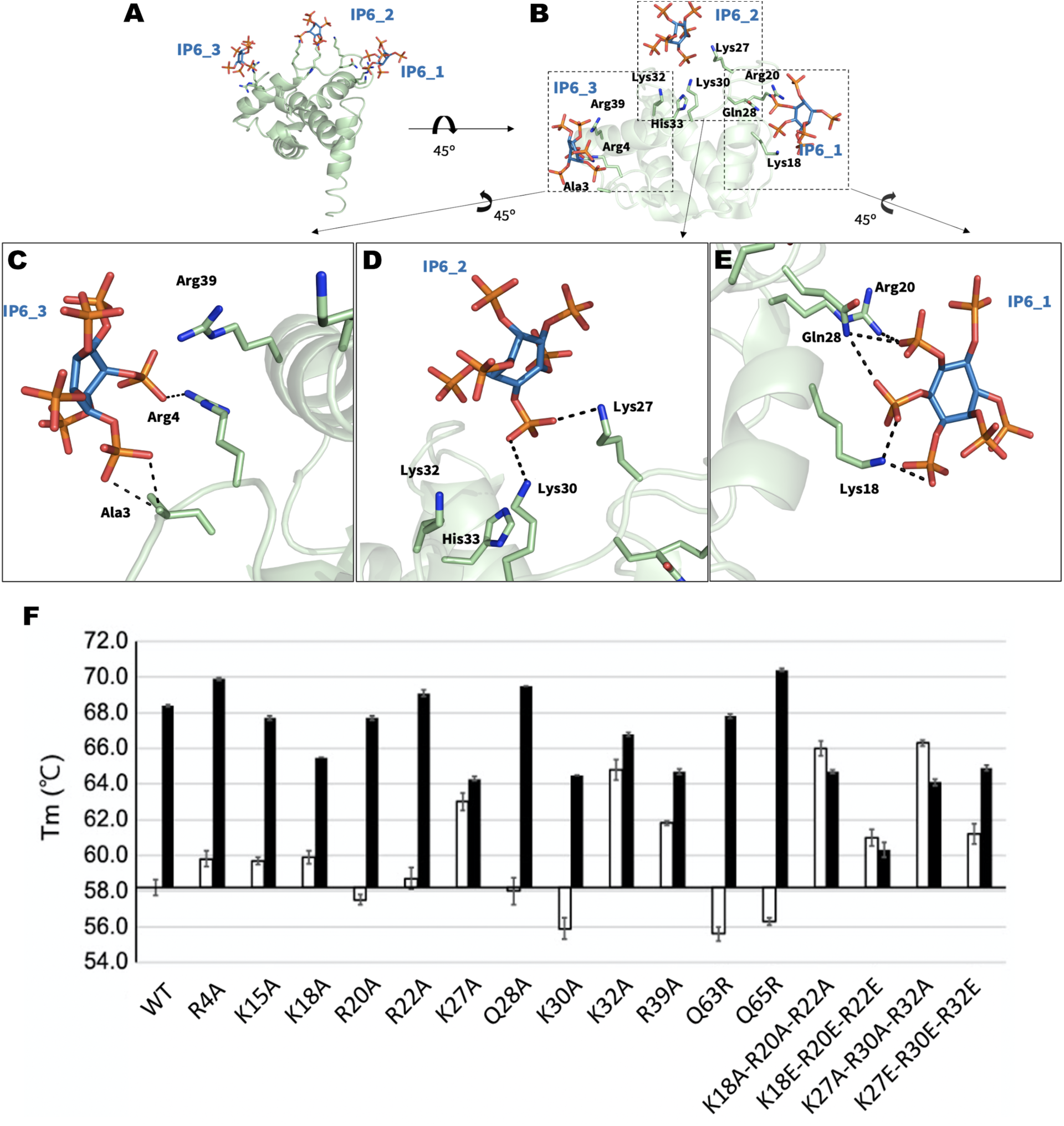
IP6-interacting amino acid residues of MA. **a** IP6 molecules near chain A and its symmetry mates of MA are shown in sky-blue and the side chain of amino acid residues in 5.0 Å is shown by sticks (pale green). **b** The three IP6 molecules are represented with their interacting residues on chain A. The interaction between **c** IP6_3 **d** IP6_2 **e** IP6_1 and chain A of the MA domain is shown in more detail. **f** DSF assay results of point mutations of MA in the presence (black bars) and in the absence of IP6 (white bars). Melting temperatures (Tm) of the wild-type and mutant proteins are indicated after normalization with wild-type protein Tm in the absence of IP6.

### DSF Assay

Melting temperatures (Tm) of the wild-type (wt) MA domain and its mutants were determined by DSF (Fig. 5F). In addition to that, the effect of IP6 presence and absence on the wt and its mutants were observed. In the absence of IP6, R4A, K15A, R20A, R22A, Q28A and Q65R single mutations affected the Tm less than 2°C while K18A, K27A, K30A, K32A, R39A and Q63R single mutations caused the notable change. Among them, K18A, K27A, K32A and R39A mutations increased the Tm according to wt-MA whereas, K30A and Q63R mutations decreased the Tm. We preferred to use ΔTm (ΔTm: Tm difference between the presence and absence of IP6) in comparisons because there is more than one factor affecting Tm in mutations in the presence of IP6 (Supplementary Fig. 14). K18A, K27A, K30A and R39A mutations decreased the Tm of MA protein in the presence of IP6, while Q65R mutation increased it. To test these residues’ combined effects, we combined these mutations and formed new triple mutants and determined their Tm. We generated two mutation sets based on these single mutation results: K18-R20-R22 and K27-K30-K32. These amino acid sets were mutated to alanine and aspartate for each mutant combination. K18A-R20A-R22A, K18E-R20E-R22E, K27A-K30A-K32A and K27E-K30E-K32E triple mutants have higher Tm than wt-MA domain in the absence of IP6. In the presence of IP6 within the DSF reaction medium, K18A-R20A-R22A, K18E-R20E-R22E and K27A-K30A-K32A triple mutations were observed to have a decreased MA Tm than the wild type. Single-residue mutants K27A, K32A, R39A and all the multiple mutants show little to no change between the presence and absence of IP6. Based on these findings, we can deduce that the mutations on IP6 binding sites inhibit IP6 interaction with MA. Finally, we conclude that presence of IP6 enhances the thermal stability of MA protein.

### IP6 has an alternative binding site for MA non-overlapping with a PIP2 binding pocket

IP6 and PIP2 are both involved in the viral life-cycle as cofactors. Previous work suggests that IP6 has a similar function with PIP2 in the viral life-cycle (Fig. 1) ^25^. To test this, we have compared the MA binding sites for both IP6 and PIP2 molecules (Fig. 6 & Supplementary Table 4). We superposed our high-resolution X-ray structure of the MA_IP6_C2 complex with an existing PIP2 bound model (PDB ID: 2H3Q) (Fig. 6C, 6E & 6F). As a result, a comparison of MA_IP6_C2 structure with MA_PIP2 complexes indicated the different binding positions for PIP2 and IP6 within the HBR. For instance, PIP2 interacts with residues Arg22, Lys27, Arg76 (PDB ID: 2H3Q) (Fig. 6C); and Arg22, Arg76 (PDB ID: 2H3Z) (Fig. 6D) respectively. Whereas IP6_1 interacts with Lys18, Arg20 and Gln28 through the hydrogen bonds in our MA_IP2_C2 structure. Furthermore, symmetry mates of chain A of MA showed that IP6 (IP6_2 and IP6_3) can bind to MA with different residues within the HBR, leading to a potential competition as a cofactor between IP6 and PIP2 (Fig. 5). One of the residues Lys27 which interacts with both IP6 and PIP2 with Myr presence, this competition may play an important role during the envelope incorporation process for making equilibrium between MA assembly on the membrane by IP6 and anchoring by PIP2-Myr interactions. In addition, the different conformation states on the C-terminus were observed between MA_IP6_C2 and MA_IP6_R32 (Supplementary Fig. 4 & 5). Our structures demonstrate that IP6 binds to MA from the nearby region but not the same amino acid residues involved in PIP2 binding (Fig. 6). These observations collectively suggest that both IP6 and PIP2 can simultaneously bind the MA. This intricate interplay of these binding events of these two host molecules may orchestrate the membrane binding and assembly either simultaneously or sequentially.

**Fig. 6:**
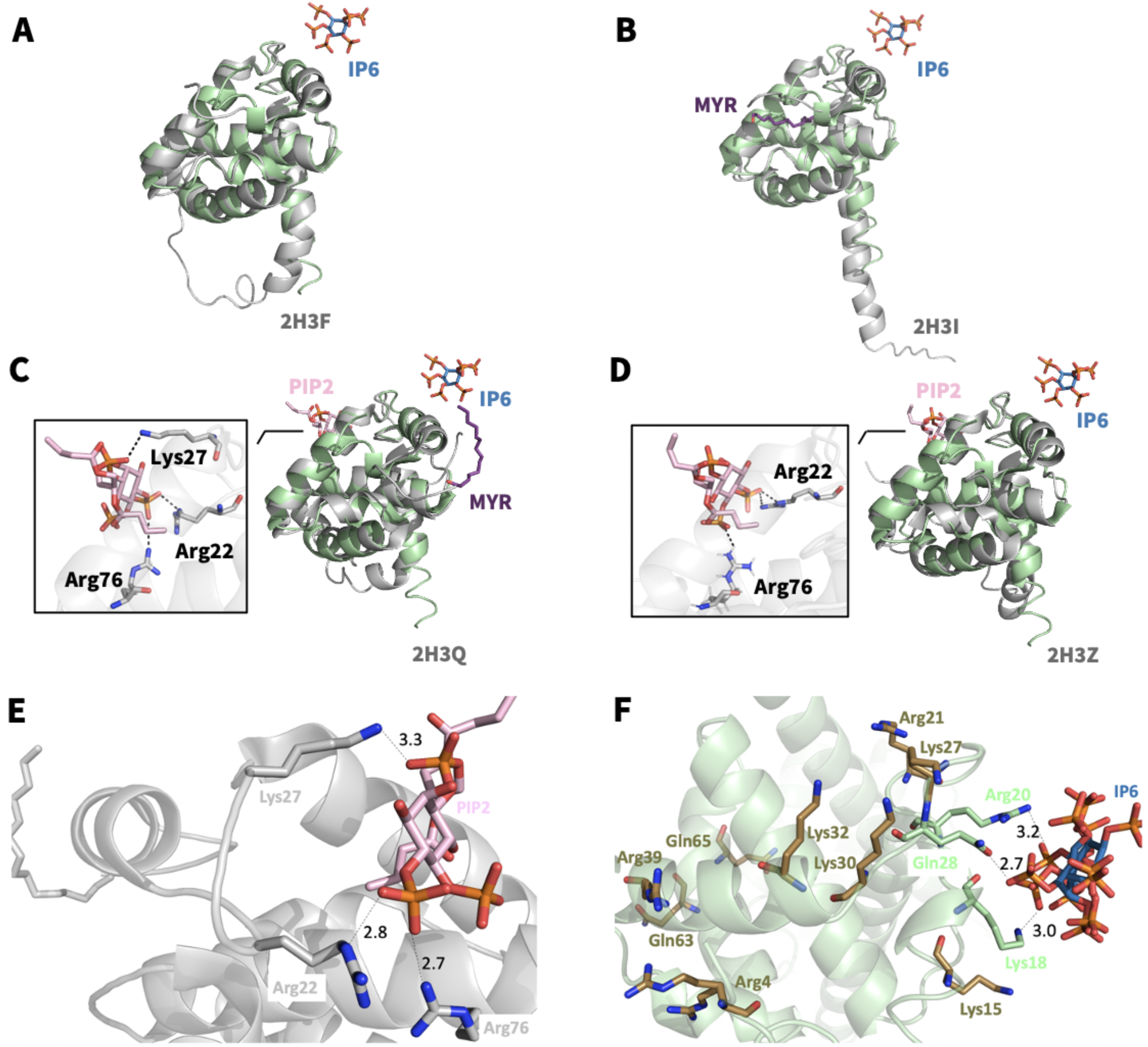
Structural comparison of MA_IP6_C2 with other MA complexes. Chain A of MA_IP6_C2 complex in green is superposed **a** with MA structure in gray (PDB ID: 2H3F), RMSD of 0.864 Å; **b** with MA structure that has myristoylation (purple) in gray (PDB ID: 2H3I), RMSD of 0.737 Å; **c** with MA structure that has myristoylation (purple) and PIP2 in gray (PDB ID: 2H3Q), RMSD of 0.742 Å; **d** with MA structure that has PIP2 in gray (PDB ID: 2H3Z), RMSD of 0.818 Å. The interaction between **e** IP6 and MA domain, **f** PIP2 and MA domain (PDB ID:2H3Q) is represented in more detail to equate the position of the corresponding molecules and angles more clearly. PIP2 is shown in color ‘light pink’, IP6 is shown in ‘sky-blue’, bound residues in ‘deep-olive’, rest of the highly basic region in ‘forest’, rest of the domain in ‘Gray’. Also, Lys27, Lys30, Lys32 and Arg39, Gln63 and Gln65 are shown to specify the regions mutated in DSF studies regarding the melting temperature. O is coded with red color, N with blue and P with orange. This figure shows that both PIP2 and IP6 bind to the MA domain within the highly basic region, however, the residues are different. Distances (within hydrogen bond boundaries) are also shown here in Angstrom. For IP6 binding, only the shortest distance between Gln28 and the IP6 is shown among 3 for clarity.

### Dynamic causal interrelations in the assembly of MA

We explored time-dependent structural fluctuations and dynamic causal residue interrelations in the hierarchical assembly of HIV-1 MA domain through its monomeric, trimeric, hexameric and dodecameric structures by utilizing their global dynamic modes. GNM based transfer entropy calculations disclosed an interplay of dynamic allosteric interactions among IP6, PIP2, membrane (envelope protein and myristoyl) and trimeric and higher oligomeric interaction sites, presented in Supplementary Fig. 6-10 and detailed in **Supplementary Results**, suggesting a dynamic order in oligomerization of MA domain.

### Order of events predicted by GNM-based transfer entropy calculations

Average ten slowest GNM modes of motion extracted from monomeric, trimeric and higher oligomer topologies were used to identify predominant directional communication paths, while average two to ten slowest dynamic modes were used to disclose subtler interaction paths. Fig. 8 features a plausible order of events as follows. **1-3.** IP6 binding on sites Lys18, Arg20 and Gln28 of monomer facilitates trimerization, possibly via trimerization site I (42-48) first, since its entropy receiving role from the IP6 binding sites is embedded in the slowest mode. For trimerization site II (64-72), similar facilitation by IP6 (Arg20) appears when we remove the slowest mode. The occupation of either trimerization sites might dynamically affect each other since there is an evident information entropy transfer between these two sites of monomer. On the other hand, PIP2 binding might trigger interactions with the envelope protein initially via Arg76 residue, a prominent entropy source to the envelope interaction sites (Leu12, Glu16 and Leu30) in the presence of the slowest mode. As Arg76 being farther than IP6 site, it can potentially generate a unique PIP2-specific response in PIP2 presence. IP6 binding and trimerization status of monomer at trimerization site II (bound to another monomer or not) might also affect envelope protein interactions with the entropy transfer from the first two sites to the latter in the presence of the slowest mode. **4.** The interaction status with envelope protein, hence membrane, affects further oligomerization. Envelope interaction sites drive the movement of oligomerization sites Arg20, Lys27, Gln28 and Lys30 overlapping with IP6 sites in average ten slowest modes of the trimer. This might indicate that the presence of membrane affects the oligomerization in high complexity (as hexamer or higher forms). **5 & 6.** PIP2 via Arg76 and trimerization sites might have a role in higher state oligomerization since these sites drive the movement of oligomerization sites in the slowest mode of the hexamer. On the other hand, oligomerization state (bound to another trimer or not) and IP6 and PIP2 binding might also affect trimer stability, since higher oligomerization sites (Arg20, Lys27, Gln28 and Lys30) also transfer entropy to trimerization sites in average two to ten slow modes. This bidirectional communication might ensure the organization of higher-order assemblies with optimum parameters such as radius of curvature to enable the formation of proper capsids eventually. **After 6.** More IP6 binding than one-per-monomer might affect further oligomerization and envelope/membrane binding. Extra IP6 binding sites (for IP6_2 and IP6_3) were seen in higher oligomerization states affecting oligomerization sites and envelope protein binding sites. Additionally, interactions among monomers in trimer via trimerization sites affect the further oligomerization possibly via adjusting the trimer components (assuming flexible interactions instead of a rigid behavior for trimers) in response to the need of relevant higher-order oligomerization state.

**Fig. 7:**
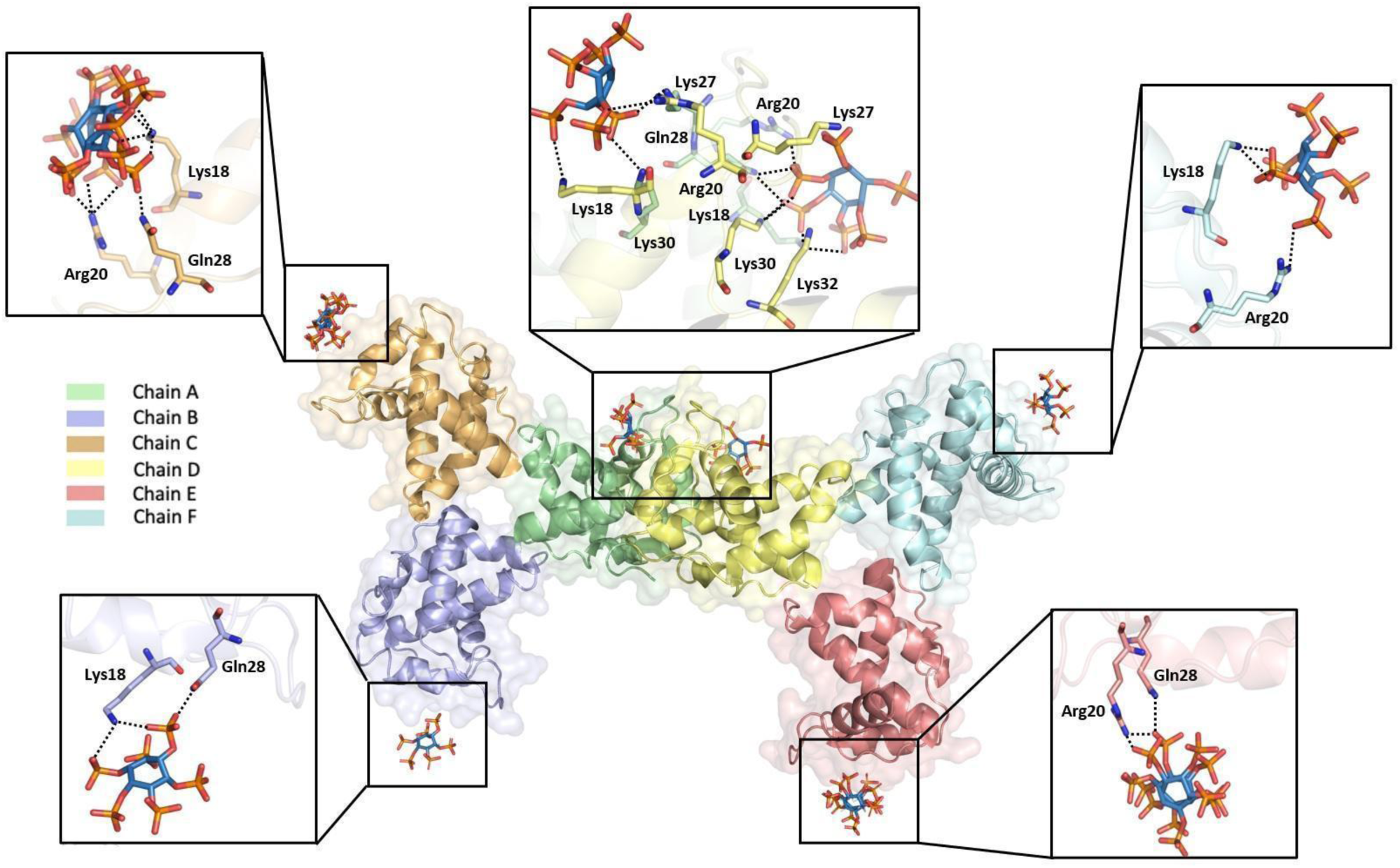
Representation of interactions on MA_IP6_C2 structure. The oligomeric structure of MA carries several IP6 molecules that interact with different residues between two trimeric proteins according to single trimer and IP6 interactions. Each chain is colored by individual colors as indicated previously. Carbon, oxygen and phosphorus atoms of IP6 are colored by sky-blue red and orange, respectively. The polar contacts between IP6 molecules and residues of MA oligomer are shown with dotted lines.

**Fig. 8:**
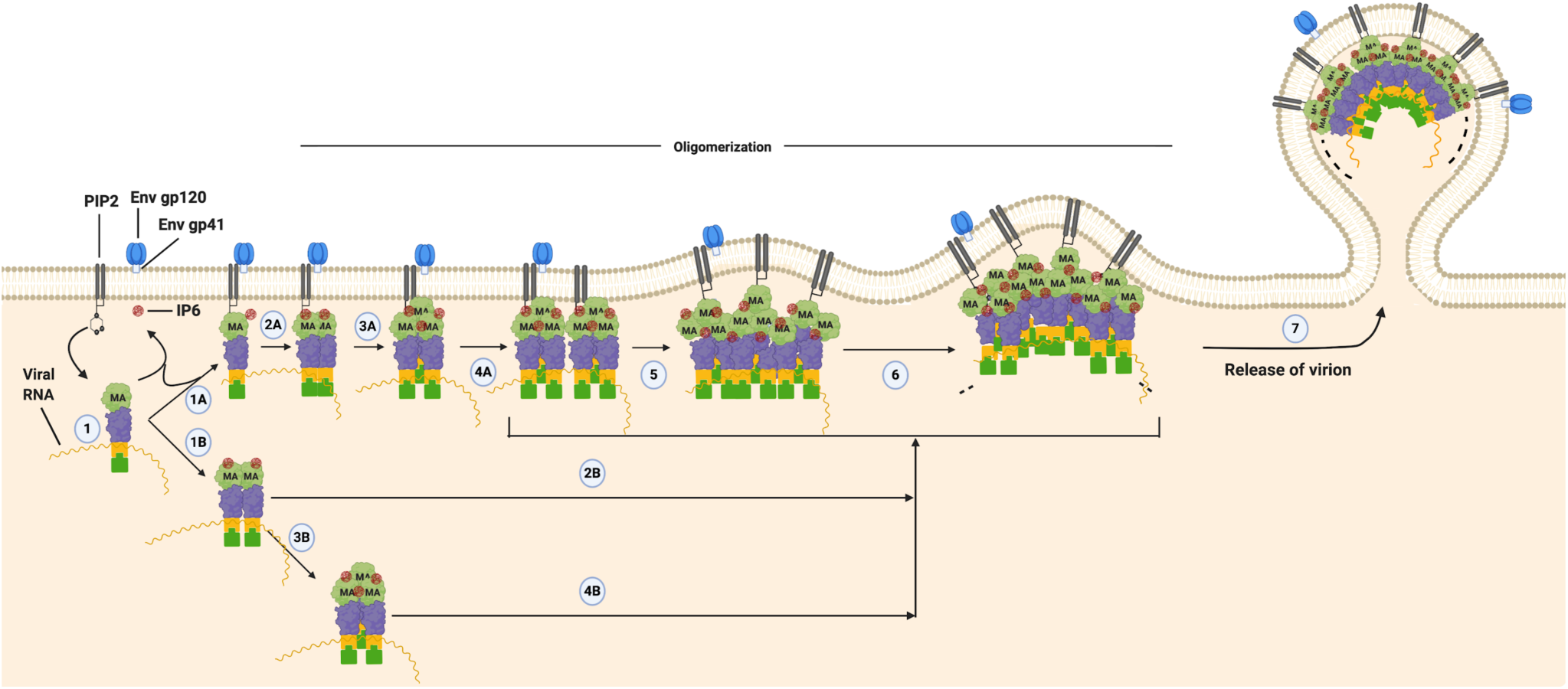
A plausible order of events for Gag oligomerization. MA domain of Gag interacts with PIP2 and IP6 to orchestrate Gag integration into the host plasma membrane (1) The interaction process is shown in the figure by labeling as A and B to represent plausible alternate kinetic pathways. According to this model, the process can be proceeded by dimerization or trimerization of the monomer form of Gag (1B & 3B, respectively). The formed dimer or trimer structure in the cytosol can be involved in the oligomerization process by the contribution of PIP2 and IP6 (2B&4B) and the progression of the higher-order oligomerization can continue in the membrane (5, 6 & 7). Besides, direct binding of the monomer form of Gag to the membrane (1A) may lead to further oligomerization (2A-4A) for the integration of Gag. IP6 binding (one-per-monomer) facilitates trimerization, possibly via trimerization sites, affecting further oligomerization and envelope/membrane binding through Env gp41 penetration to the cellular membrane. Together with this, PIP2 binding might trigger interactions with the envelope protein, while IP6 binding and the status of trimerization may affect membrane interactions. Oligomerization sites with IP6 and PIP2 binding might also affect trimer stability; this distinct bidirectional causality along with membrane and myristoyl interactions likely ensures the high order assemblies with optimum parameters.

## DISCUSSION

Different HIV-1 proteins, enzymes and metabolic steps from the first contact with the host to the production of new viral particles have crucial roles in the HIV-1 life cycle and have been potential drug targets. Pr55^Gag^ protein regulates HIV-1 assembly, budding and maturation of virions. Working with this essential protein and its complexes may provide a further understanding of the formation of virions ^4^. Pr55^Gag^ contains MA, CA, NC, and p6 domains, where the MA domain regulates Pr55^Gag^ membrane binding and assembly ^49, 50^. Given the key importance of the MA domain in the virion cycle, inhibition of this protein may provide an effective treatment against HIV and makes it an important drug target ^18, 51^.

Previous studies revealed the physical interaction between both PIP2-MA and IP6-MA *in vitro* ^10, 25, 52–55^. It is shown that inositol phosphates are crucial cofactors involved in the complex HIV-1 assembly process ^7, 25^. The need for the presence of IP6 in the viruses, even in “encapsidation” alone, is a milestone for HIV virion formation ^22, 42^. On the other hand, IP6 has a strong influence on not only CA but also MA trimerization and has a key role in assembly and encapsidation ^25^. Additionally, IP6 has been shown to interact with MA at the highest level among inositol phosphates species ^51^. The requirement of IP6 in the different steps of the viral life-cycle makes IP6 the target of the study. Therefore, any high-resolution structural information about the IP6-MA complex will provide invaluable details of the assembly process. For instance, our study demonstrates that IP6 and PIP2 binding is not mutually exclusive and compatible with taking place together for localization and conformation of MA on the plasma membrane. Additionally, the C-terminus of the MA domain, which is crucial for CA domain binding and conformation of Gag, was determined with different conformation states by different oligomerization structures such as trimer and hexamer (Supplementary Fig. 4). The different conformations of C-terminal by trimer or hexamer MA with IP6 interaction can induce an overall conformational change of Gag protein assembly by squeezing and bending the proteins’ opposite terminals. Our hexamer structure with IP6 binding displays these possible conformations (Supplementary Fig. 11).

Our high-resolution MA_IP6_C2 structure demonstrates critical interactions of IP6 with MA explains how IP6 may favor trimeric state (Fig. 7). It may also provide an explanation for the previous observation that the presence of IP6 favors the MA protein’s higher-order oligomeric state of its trimers ^25^. These interactions are largely mediated through H-bonds by the basic side chain residues Lys and Arg together with polar uncharged Gln residues. Importantly, two nearby IP6 molecules are present at the site where the two MA trimers form the hexamer are involved in pivotal intermolecular interactions. For the selected trimer-trimer interaction site based on crystal packing and contact patterns, the interaction between IP6 and MA are mediated by the chain A residues Arg20, Lys27, Gln28, Lys30 and the chain D residues Lys18, Arg20, Lys30, Lys32 (Fig. 7). Lys27 carries specific importance for the comparison of IP6 and PIP2 binding because this residue interacts with both of these molecules, leading a plausible mechanism between PIP2 and IP6 (Fig. 6 & 7). This result suggests that after the induction of Gag oligomerization, IP6 attach to this region may replace it with PIP2 when Gag reaches the membrane; and IP6 binding helps the MA for generating higher-order oligomerization, showing that IP6 and PIP2 binding may be compatible. Besides, chain B, C, E and F form interactions with IP6 molecules by Lys18, Arg20 and Gln28 residues in varying binding angles, distances and conformations (Fig. 7). This result may show that H-bond interactions gave specificity, but electrostatic interaction of side chains resulted in different angles, distances, and conformations for IP6 and MA binding in each of the trimers. The main distinction of the MA trimer dimerization interface is the involvement of three different amino acid interactions at Lys27, Lys30 and Lys32 besides monomer level MA and IP6 interactions. We have noticed that the residues mutated in DSF analysis (Fig. 5F) are not directly related to monomeric level binding of MA to IP6; therefore, they either have a role in oligomerization (likely Lys27, Lys30 and Lys32) or structural stability of MA domain (Gln63 and Gln65). Another possible explanation for this situation is that there is more than one location where the IP6 can bind and interact with the MA domain, each site responsible for different kinetic pathways in the assembly process (Fig. 5 & 7).

Our data explain the previous *in vitro* results in which they showed that in the presence of IP6, MA enhances the formation of trimers ^25^. Acidic IP6 molecules occupy corners of the MA protein promoting the protein trimerization from the sides by attracting their basic charges (Supplementary Fig. 12 & Supplementary Fig. 13). Trimerization of the MA may provide more stability and binding of IP6, thereby increasing the Tm of the MA protein. The IP6 molecules present in our structures allowed us to provide a model for high-order oligomerization of MA and suggest a role for IP6 in these multistep structural rearrangement events (Supplementary Fig. 11). It was shown that not delivery of Gag to the membrane but virion assembly during budding is perturbed by reduced cellular IP6 levels ^22^. This may suggest that MA-IP6 complex occurs during virion assembly and budding. Here, we suggest a new model for the formation of higher-degree MA oligomerization induced and stabilized by IP6 binding. IP6 binding stabilizes the expanded conformations of an oligomeric and more organized state on the plasma membrane (Supplementary Fig. 11B-D). Although the role of IP6 on budding mechanisms is still unclear, it is possible that the interactions of PIP2 and IP6 binding can occur simultaneously in this mechanism, since they have non-overlapping binding sites on MA.

To explain the effect of IP6 on MA protein, its membrane interaction and higher-order oligomerization, causal dynamic interactions in different oligomerization states of MA were explored by using transfer entropy TE calculations for the directional allosteric interactions (Supplementary Fig. 6-10). While some residues convey information as entropic sources, some receive it as entropic sinks. With their interchangeable roles as entropic sources and sinks, IP6/PIP2 binding sites and trimeric interfaces of MA display a dynamic interplay among each other, resulting in a bidirectional information flow manifested with the dissection of slow modes of motion/informational entropy. In trimer, the interactions sites with envelope protein and myristoyl molecules are the major entropic sources. The dynamic fluctuations of these sites will possibly lead to the conformational and/or dynamic changes in the various regions of the trimer including trimerization sites along with IP6 and PIP2 binding sites. At this early oligomerization step as a trimer, envelope protein and myristoyl interaction sites might be more important due to the membrane attachment, relaying information regarding the membrane presence toward the trimerization regions along with IP6 and PIP2 binding sites. On the other hand, a similar bidirectional causality among IP6 and PIP2 sites and trimeric interfaces (interfaces of the MA domains) are observed in both structures of the hexamers despite their distinct structural organizations. Here, PIP2 and trimeric interfaces appear as the main drivers (i.e., entropic sources) with IP6 causing subtler allosteric interactions. In conclusion, allosteric signals are bidirectional, and the dynamic information is conveyed in opposite directions by recruiting different modes of motion. These movements form a dynamic repertoire for the protein. It provides functional conformation and dynamical changes in response to the perturbations and environment by exerting certain relevant modes of motion. Those results verified the previous findings of IP6 effect on MA trimerization; PIP2 and myristoyl interactions on membrane attachment supported the concepts of IP6 on higher-order oligomerization; and alteration of membrane localization and binding with PIP2. Still, further molecular dynamics studies must be carried on for higher degree oligomerization of MA in the presence of IP6. These studies can be divided into two subtopics: (i) trimer dimerization; (ii) simulation of dissecting different binding modes of IP6 on MA and its effect on the stability of the larger oligomeric state.

Dynamic interaction of PIP2 and IP6 on MA is crucial for promoting Gag localization to the plasma membrane during the viral maturation. That is why, targeting MA through these two small molecules carries significant importance for molecular design of new generation anti-HIV-1 agents to block membrane localization of Pr55^Gag^ and virion formation. The data presented here added a new structural perspective to the predicted interaction between MA-IP6 for further studies. Our suggested model reveals potential new target sites to develop novel therapies for the treatment of HIV-AIDS.

## Accession codes

Coordinates and structure factors have been deposited in the Protein Data Bank under accession 7E1K, 7E1J and 7E1I for data sets MA_IP6_R32, MA_IP6_C2 and MA_IP6_SFX respectively.

## Author Contributions

HD, MF, TS and MO designed the project. HIC, HT, KK and RK prepared the samples for cryo and SFX studies. RGS and HD build the coMESH injectors. CHY and ZS executed the SFX data reduction and provided realtime XFEL data analysis. KK, FY and HD refined the cryo and SFX structures. The experiments were executed by HIC, HT, KK, RK, KA, KM, RGS, ZS, ML. GNM-based transfer entropy calculations were performed by BA and TH. Data were analyzed by HD, TS, MO, MF, TH, BA and RGS. The manuscript was prepared by HD, HIC, ED, BY, EA, GY, MY, FBE, AS, OG, SOB, TH, MO, MF and TS with input from all the coauthors.

## Acknowledgements

HD acknowledges support from the National Science Foundation (NSF) Science and Technology Centers grant NSF-1231306 (Biology with X-ray Lasers, BioXFEL). This publication has been produced benefiting from the 2232 International Fellowship for Outstanding Researchers Program of TÜBİTAK (Project No: 118C270). However, the entire responsibility of the publication belongs to the owner of the publication. The financial support received from TÜBİTAK does not mean that the content of the publication is approved in a scientific sense by TÜBİTAK. HD would like to thank Michelle Young, Ritu Khurana, Lori Anne Love and Tracy Chou for their invaluable support and discussions. The LCLS is acknowledged for beam time access under experiment no. cxils9717. Use of the Linac Coherent Light Source (LCLS), SLAC National Accelerator Laboratory, is supported by the U.S. Department of Energy, Office of Science, Office of Basic Energy Sciences under Contract No. DE-AC02-76SF00515. This work was supported in part by Grant-in-Aid for Scientific Research (C) (17K08861), Grant-in-Aid for Research Activity Start-up (19K23802) and Grant-in-Aid for Scientific Research (B) (20H03365) from the Japan Society for the Promotion of Science (JSPS), Grant-in and Aid for JSPS Research Fellow (17J11657), and the Platform for Drug Discovery, Informatics, and Structural Life Science (Ministry of Education, Culture, Sports, Science and Technology (MEXT), Japan Agency for Medical Research and Development (AMED)) and Basis for Supporting Innovative Drug Discovery and Life Science Research from Japan Agency for Medical Research and Development (AMED) under Grant Number JP20am0101071 (support number 2029). TH acknowledges TUBITAK (Project No: 115M418).

## Supplementary Results

### Dynamic causal interrelations in monomeric, trimeric, hexameric and dodecameric structures of MA domain

#### Monomer (MA_IP6_C2)

Supplementary Figure S6 displays the causal interrelation of residue pairs in average ten (Figure S6A) and average two to ten (Figure S6B) slowest dynamic modes of monomer, pointing to interactions among IP6 and PIP2 binding sites, trimerization sites I & II and envelope and myristyl interaction sites. The casual interrelations are rather promiscuous, yet some distinct causal interrelations are still observed as follows.

IP6 might be the major player in the monomer state of MA since topology imposes as entropy sources the recruit of all three IP6 binding sites Lys18, Arg20 and Gln28 in the most global mode as well as Arg20 in average two to ten slowest dynamic modes. IP6 binding may facilitate trimerization possibly via trimerization sites I (42-48) and II (64-72) as appear on nTE maps calculated with and without the slowest dynamic mode, respectively. The occupation of either trimerization site might also dynamically affect each other as shown by information transfer between the two. IP6 binding and trimerization status (bound to another monomer or not) at trimerization site II might also affect envelope protein (Alfadhli, 2016) and myristoyl (Saad, 2007) interactions by influencing the dynamics of responsible sites. For illustration, the causal interrelations of Lys18 (Supplementary Figure S6A) and Arg20 (Supplementary Figure S6B) of IP6 binding sites were shown on the monomer (on the right).

On the other hand, different PIP2 sites transfer information via using distinct dynamic modes (Arg76 and Arg22 in average ten and two to ten slowest dynamic modes, respectively; Supplementary Figure S6) to entropy receivers/sinks indicating a more delicate communication for PIP2. PIP2 binding via Arg76 might trigger interactions predominantly with envelope protein while also affecting myristoyl interaction sites. This likely generates a unique PIP2 specific response by recruiting a residue farther than the IP6 site.

#### Trimer (MA_IP6_C2)

Supplementary Figure S7 displays the causal interrelation of residue pairs in average ten slowest dynamic modes of the trimer. While all constituent chains behave similar to each other, entropy sources cluster in the vicinity of IP6 binding sites on these chains (yet shown only on chain A for simplicity). Among the entropy sources, envelope protein (Leu13, Glu17, Leu31, Val35 and Glu99) and myristoyl (Val7, Ile34, Val35, Leu51 and Glu52) interaction sites were both seen to drive the movement of trimerization sites I (Glu42-Pro48) and II (Leu64-Ser72) along with IP6 (Arg20, Gln28) and PIP2 (Arg22, Lys27 and Arg76) binding sites (also overlapping with higher oligomerization sites Arg20, Lys27, Gln28 and Lys30 than trimeric form). These affected entropy receivers are shown on chains B and C with red arrows. As exemplified on the right of Figure S7, the causal effects of envelope-protein-interacting residue Leu13 (chain A) are distinctive as shown particularly at trimerization sites I and II. This implies that the interaction status with envelope protein in collaboration with IP6 binding might trigger the oligomerization in high complexity. When the slowest dynamic mode is removed, a rather promiscuous causal interrelation appears (not shown). This might emphasize the role of the slowest dynamic mode in directing distinct causal signals from envelope and myristoyl interaction sites toward the IP6 and PIP2 binding sites along with both trimerization sites.

Additionally, if only five slowest instead of ten slowest dynamic modes are considered, the entropy sources would include also IP6 binding sites (Lys18 and Arg20) in addition to nearby sites (data not shown); overall indicating an important dynamic role for IP6 binding or nearby residues in initiating and facilitating the trimeric and higher oligomeric assembly process.

#### Hexamer (MA_IP6_C2)

Supplementary Figure S8A displays the causal interrelation for each residue pair in average of ten slowest dynamic modes of the hexamer. The outmost chains C and F work as strong entropy sources in general, followed by chains B and E. On the other hand, IP6 binding sites of chains A and D overlapping with the higher oligomerization sites are responsible for trimer dimerization, mostly acting as entropy sinks and receiving the dynamic signal. Specifically, PIP2 (Arg76) binding site and trimerization sites I (Glu42-Pro48) and II (Leu64-Ser72) are the main entropic sources on the hexamer (indicated by black arrows only on trimer with chains A, B, C for simplicity), which drive the movement of IP6 binding sites and their C-terminal neighbors (Lys18, Arg20, Gln28 and Asp93-Asp96 in black ellipses, indicated by the red arrow on chains D and E). As exemplary cases, the causal effects of Ala45 and Arg76 (chain B) clearly demonstrate the dynamic driving capacity of these residues on the movements of IP6 binding and nearby residues in the hexamer.

Supplementary Figure S8B, on the other hand, shows the causal interrelation for each residue pair in average two to ten slowest dynamic modes of the hexamer. IP6 and PIP2 binding sites along with myristoyl interaction sites appear as main entropy sources (indicated on chains A and B by black arrows) that drive the movement of trimerization sites I (Glu42-Pro48) and II (Leu64-Ser72), pointed by red arrows on chain A, C, E and F. The affecting residues of Lys18 and Arg22 (chain exemplify IP6 and PIP2 binding sites driving the movement of both trimerization sites. Interesting to note here is that the distinct bidirectional causality could be observed with the dissection of information entropy into different dynamic mode sets, enabling to disclose both directions of allosteric signaling between IP6 binding and trimerization sites. On the other hand, PIP2 binding sites appear as a common player in both directions of the allosteric signaling between IP6 binding, trimerization and oligomerization sites. Conclusively, the emerging interplay between distinct interface residues (of monomers and trimers) might regulate the higher-order assembly of monomers.

#### Hexamer (MA_IP6_R32)

Supplementary Figure S9A displays the causal interrelation for each residue in an average of ten slowest dynamic modes of the hexamer. While all constituent chains behave similarly to each other, PIP2 binding site Arg76 and trimerization sites I (Glu42-Pro48) and II (Leu64-Ser72) are the main entropic sources on each chain (represented by black arrows on chain A for simplicity), which drive the movement of IP6 binding regions and their C-terminal neighbors (Lys18, Arg20, Gln28 and Asp93-Asp96) along with myristoyl (Leu51, Glu52) interaction sites (pointed by red arrows on chain C for clarity). As an example, the causal effects of Ala45 and Arg76 (chain A) indicate the driving role of these residues on IP6 and nearby residues. The second figure on Supplementary Figure S9B, on the other hand, shows the causal interrelation for each residue pair in average two to ten slowest dynamic modes of the hexamer. When the slowest dynamic mode is discarded, IP6 and PIP2 (1,3) binding and myristoyl (2) interaction sites on each chain (yet shown on chain B only for simplicity) are seen to drive the movement of trimerization sites I (Glu42-Pro48) and II (Leu64-Ser72) which indicated by red arrows on chains A and E. The affected residues of Lys18 and Arg22 (chain B) exemplifies IP6 and PIP2 binding sites as driving the movement of both trimerization sites. PIP2 binding sites appear as common players in both directions of the allosteric signaling between IP6 binding and trimerization sites. These observations suggest that IP6 and PIP2 can affect each other and the protein differently. The intricate interplay of these two ligand binding events may orchestrate the membrane interactions and assembly.

The role of trimerization site dynamics is so robust in causal interactions of hexamer that even different configurations of hexamer result in a similar pattern of directional information flow orchestrated among the same regions.

#### Dodecamer (MA_IP6_P1)

Supplementary Figure S10 shows the causal interrelation for each residue pair in average ten slowest dynamic modes of dodecamer (MA_IP6_P1). As being the outmost chains, G, H and I are the main entropic sources of dodecamer. Trimerization sites I and II along with myristoyl interaction and higher state IP6 binding sites behave as driver residues on multiple chains. These sites mainly affect IP6 and PIP2 binding sites along with envelope protein interaction sites. Only on chain K (larger cumulative net transfer entropy than zero), IP6 and PIP2 binding sites act as entropy sources for the other IP6 and PIP2 binding sites on chains A, B, C. As exemplary cases, the causal interrelations of Arg4 (a higher oligomer IP6 binding residue on chain K) and Leu68 (located at trimerization site II on chain J) were shown. Without the slowest dynamic mode, chains D, E and F are the main entropic sources, followed by chains J, K, L (data not shown). With or without the slowest mode, trimerization sites I and II along with myristoyl interaction and higher state IP6 binding sites behave as driver residues on multiple chains. These sites mainly affect IP6 and PIP2 binding sites along with envelope protein interaction and trimerization sites of other trimers.

In summary, more than one IP6 binding per-monomer might affect further oligomerization and envelope/membrane binding. Trimerization sites affect further oligomerization possibly via adjusting the trimer components (assuming flexible interactions among constituent monomers instead of a rigid behavior for trimers) in response to the need for relevant higher-order oligomerization states as appear when the slowest dynamic mode is excluded.

## Supplementary Figures

**Supplementary Fig. 1:**
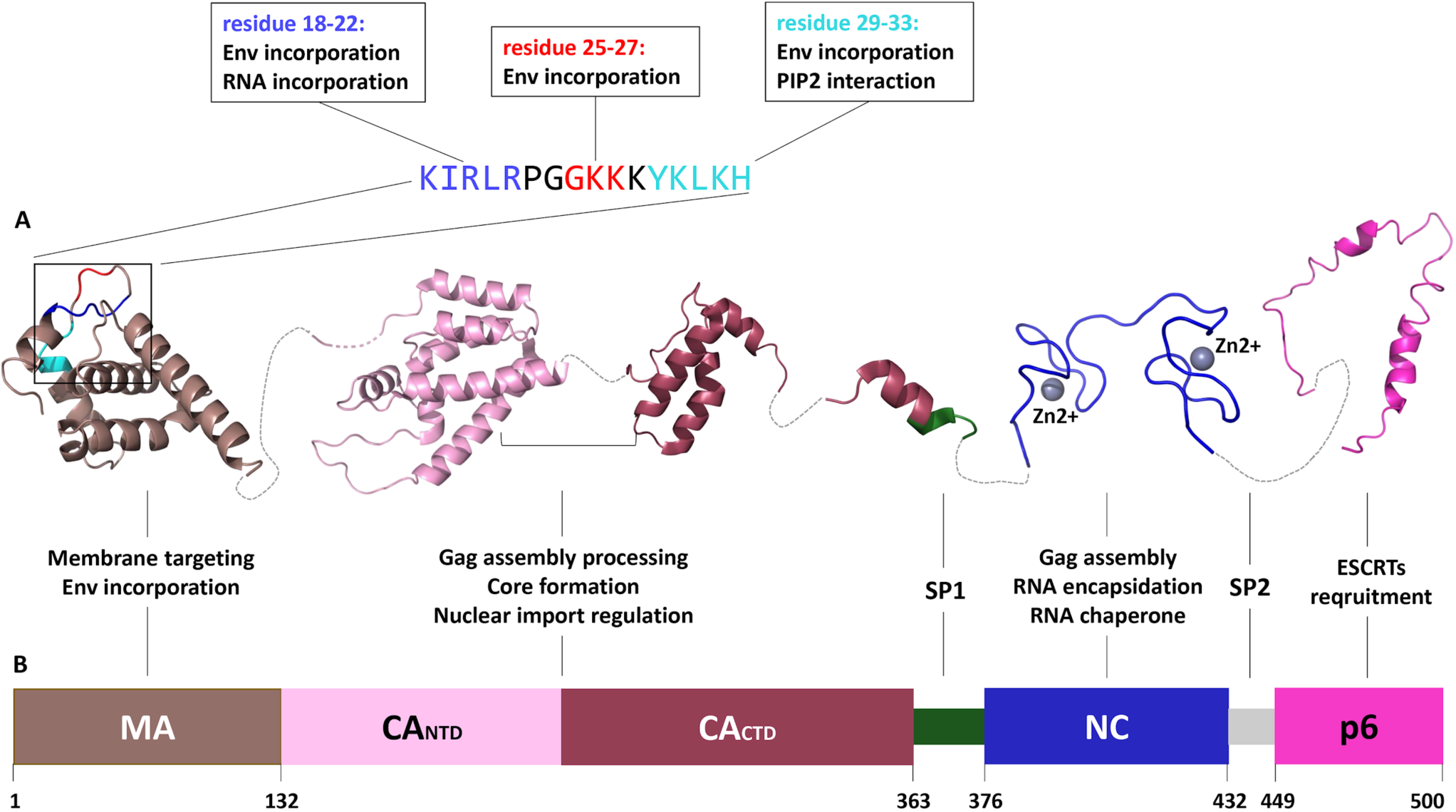
Representation of the Gag precursor polyprotein domains. **a** Structural model of the separated and extended Gag domains. The significance of the N-terminal of the MA domain is shown sequentially and its description is demonstrated in the boxes. The domains are assembled by high-resolution structures available in Protein Data Bank (PDB). Linker regions are shown with dashed lines. **b** Schematic representation of the putative Gag domain boundaries. The beginning and end of each domain are labeled as the residue number and the functions of corresponding domains are indicated.

**Supplementary Fig. 2:**
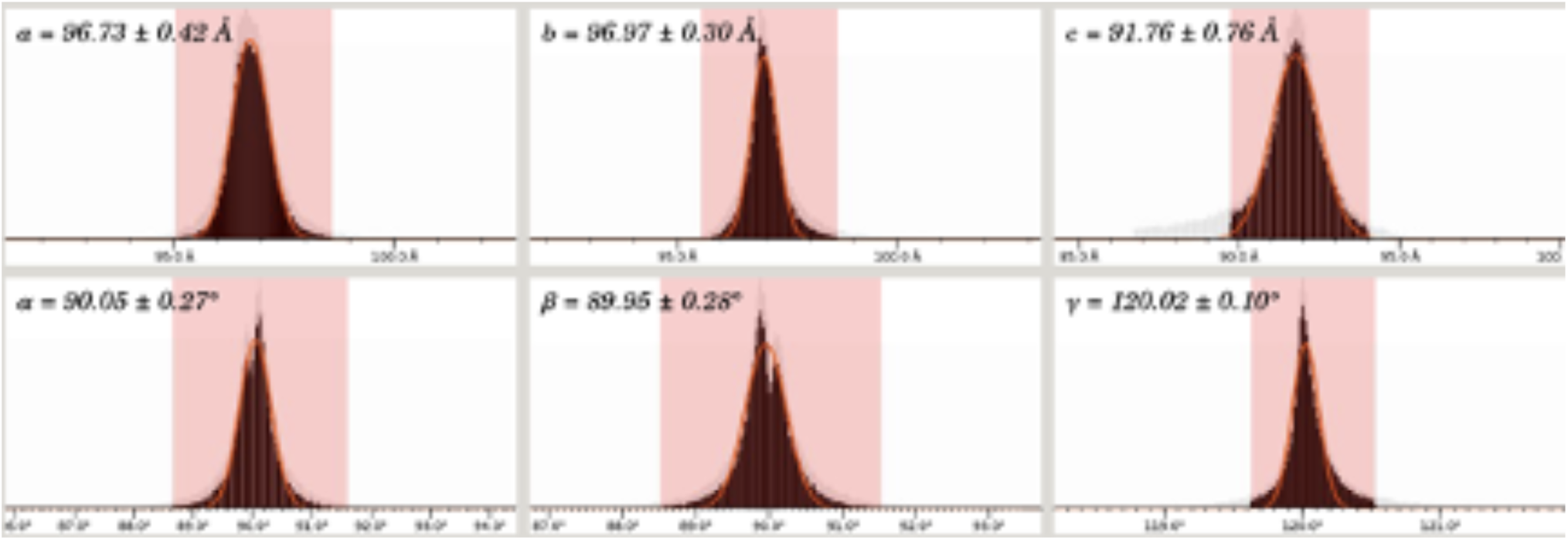
Unit cells of MA_IP6_SFX dataset. Indexing cell file generated by *CrystFEL* unit cell file version 1.0

**Supplementary Fig. 3:**
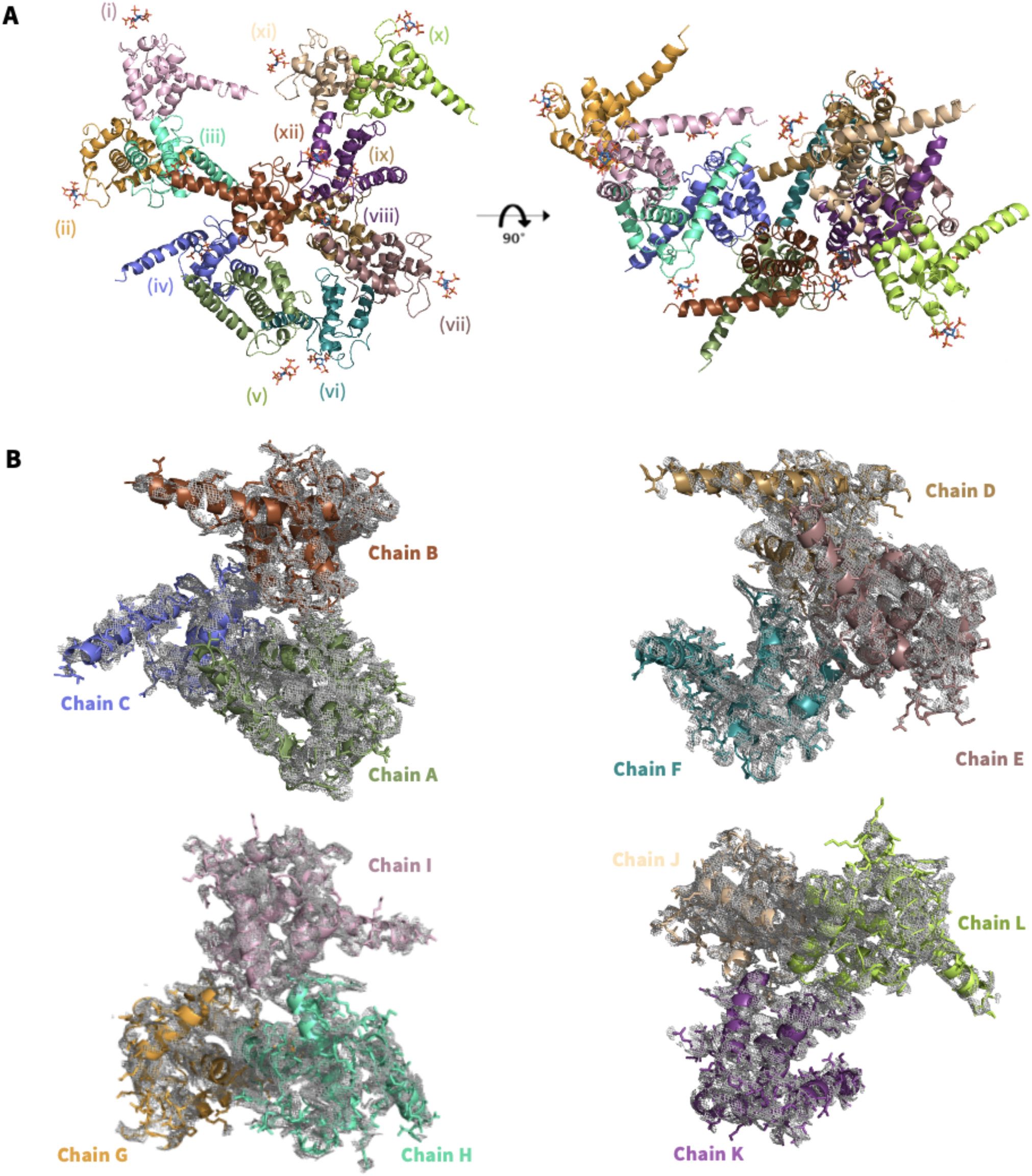
SFX structure of MA_IP6 complex. **a** The overall structure of the MA_IP6_P1 complex is colored based on each chain. There are a total of 16 molecules in the asymmetric unit cell. Carbon, oxygen and phosphorus atoms of IP6 molecules are colored by sky-blue, red and orange, respectively. **b** A 2*F*o-*F*c simulated annealing-omit map for each trimer is shown in the gray mesh.

**Supplementary Fig. 4:**
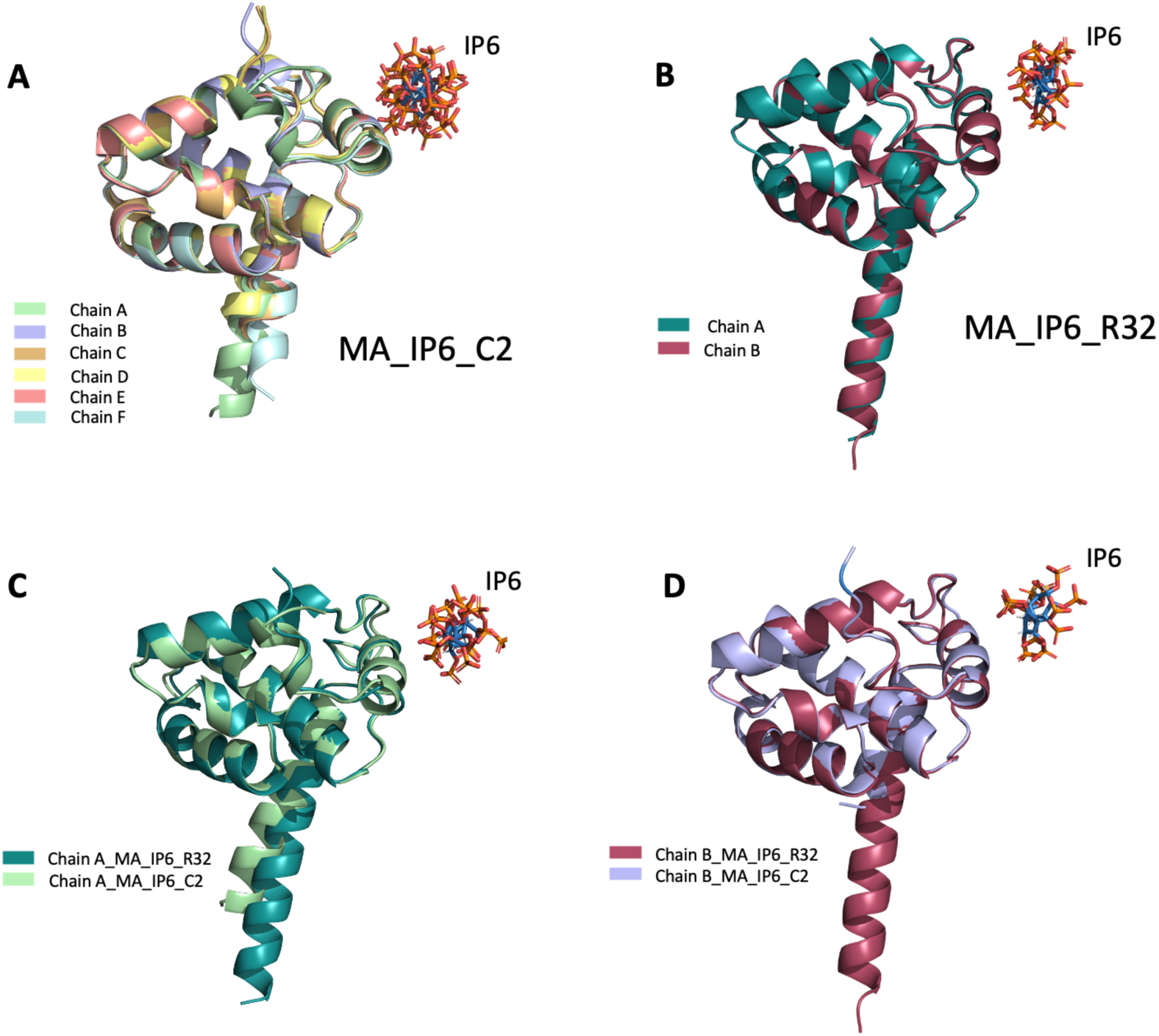
Superposition of chains for two crystal forms. **a** Each chain of MA_IP6_C2 complex is superposed with an overall RMSD of 0.286 Å. **b** Two chains of the MA_IP6_R32 complex are superposed with an RMSD of 0.200 Å. **c** Chain A of MA_IP_C2 complex is superposed with chain A of MA_IP6_R32 complex with RMSD of 0.370 Å. **d** Chain B of MA_IP_C2 complex is superposed with chain B of MA_IP6_R32 complex with RMSD of 0.317 Å.

**Supplementary Fig. 5:**
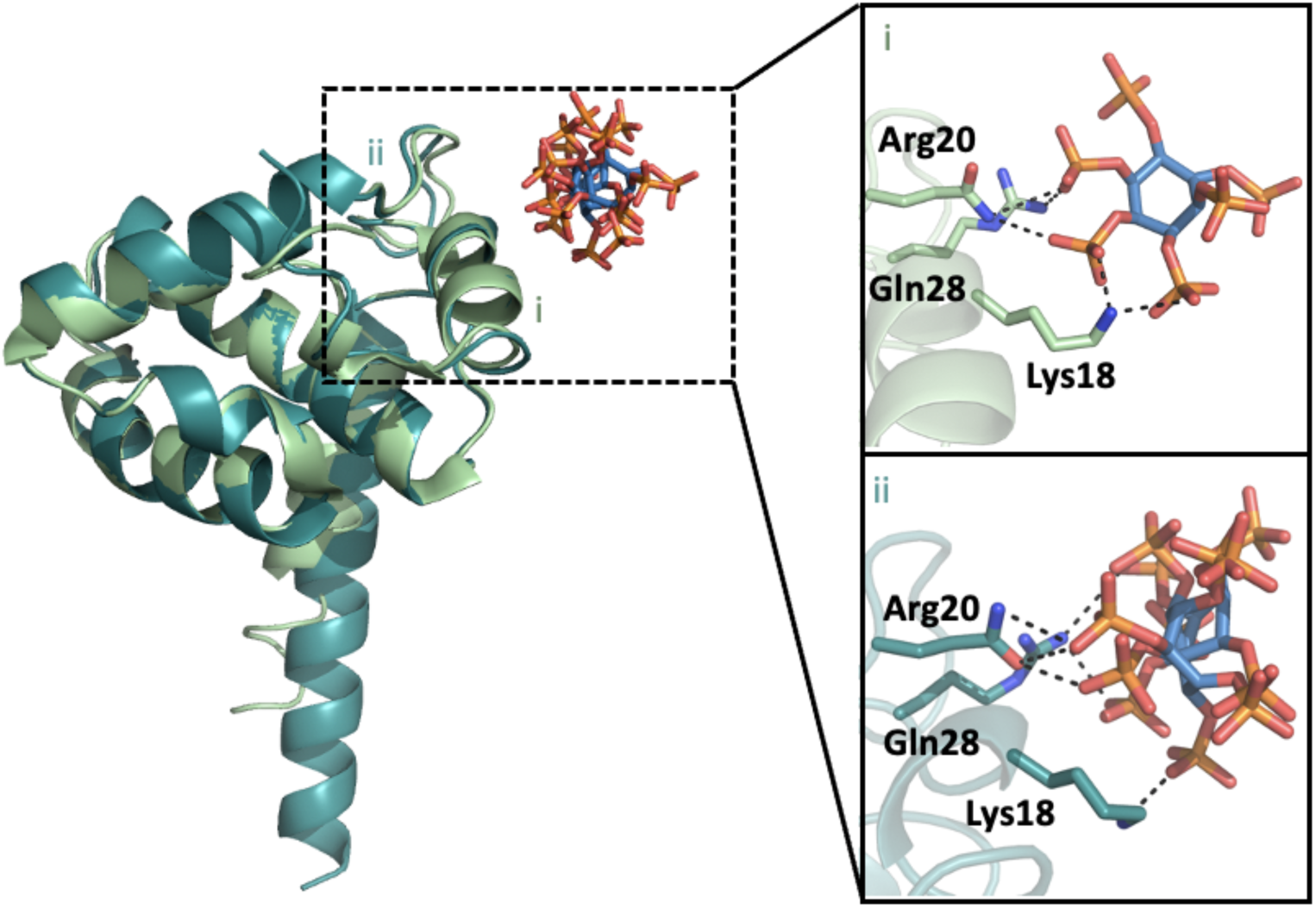
Comparison of active residues in the binding pocket for the chain A of MA_IP6_C2 (i) and MA_IP6_R32 (ii) structures. Chain A of MA_IP_C2 (Pale green) complex is superposed with chain A of MA_IP6_R32 (Deep teal) complex with RMSD of 0.370 Å. Hydrogen bonds are shown with dashed lines.

**Supplementary Fig. 6:**
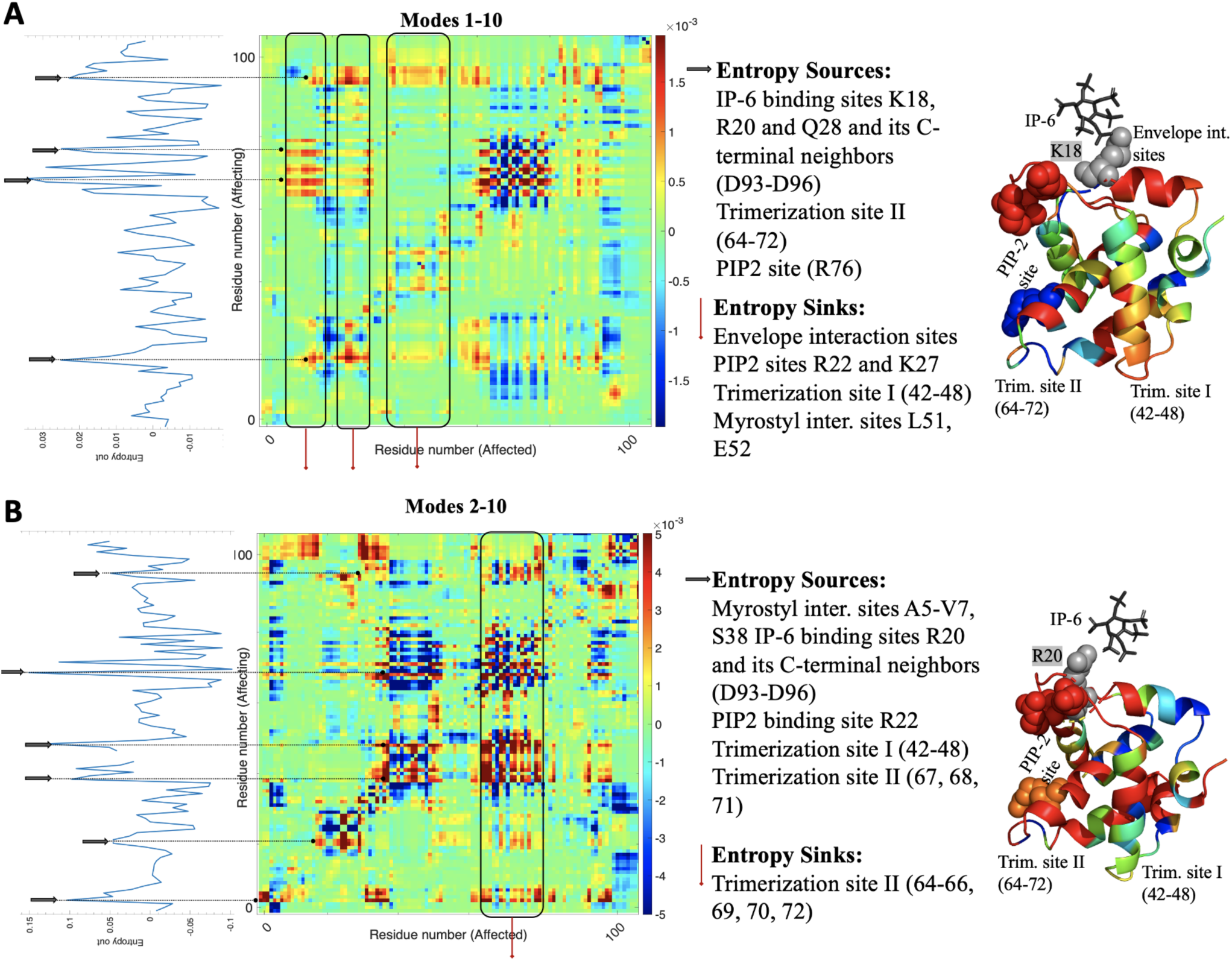
Net transfer entropy (nTE) maps of monomer (MA_IP6_C2) showing the causal interrelation for each residue pair (affecting versus affected) calculated by using an average of ten slowest modes (A) and average two to ten slowest modes (B). Cumulative transfer entropy plots (cTEs) corresponding to each nTE map are given on the left. On the right, the causal interrelation of exemplary residues from respective nTE maps, K18 **(A)** and R20 **(B)** represented with gray spheres, with the rest of residues on monomer are color-coded on structure from highest positive (red) to lowest negative (blue) nTE values of the relevant residue. Entropy sources and sinks for chain A are listed and indicated respectively with black and red arrows.

**Supplementary Fig. 7:**
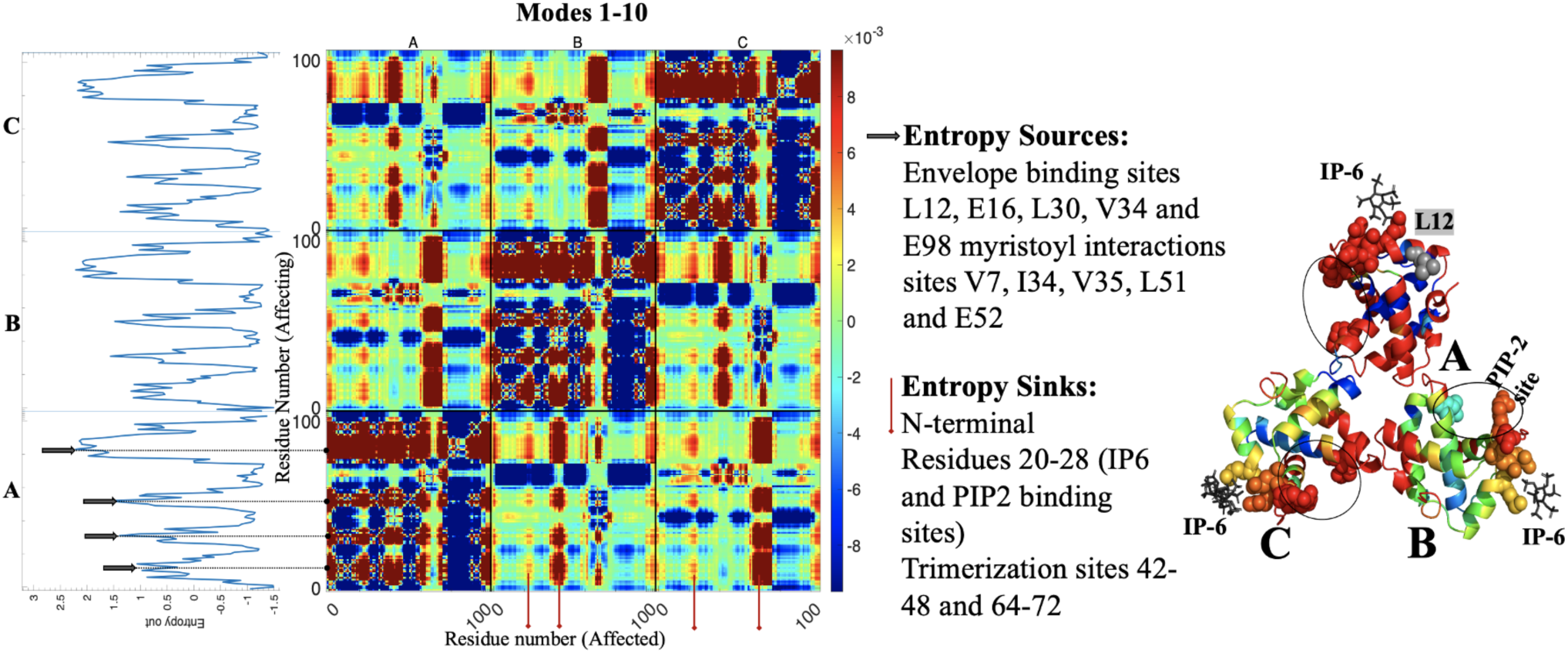
Net transfer entropy (nTE) map of the trimer (MA_IP6_C2) showing the causal interrelation for each residue pair (affecting versus affected), calculated by using an average of ten slowest modes. Cumulative transfer entropy plots (cTEs) corresponding to each nTE map are given on the left. As an exemplary case, the causal interrelation of L12 represented with gray spheres (chain A) with the rest of residues on trimer are shown, color-coded from highest positive (red) to lowest negative (blue) nTE values. The affecting and affected residues with maximum cTE values (i.e., affecting the others and affected by the others at the most) are respectively entropy sources and entropy sinks, listed and indicated with black (chain A) and red (chains B and C) arrows.

**Supplementary Fig. 8:**
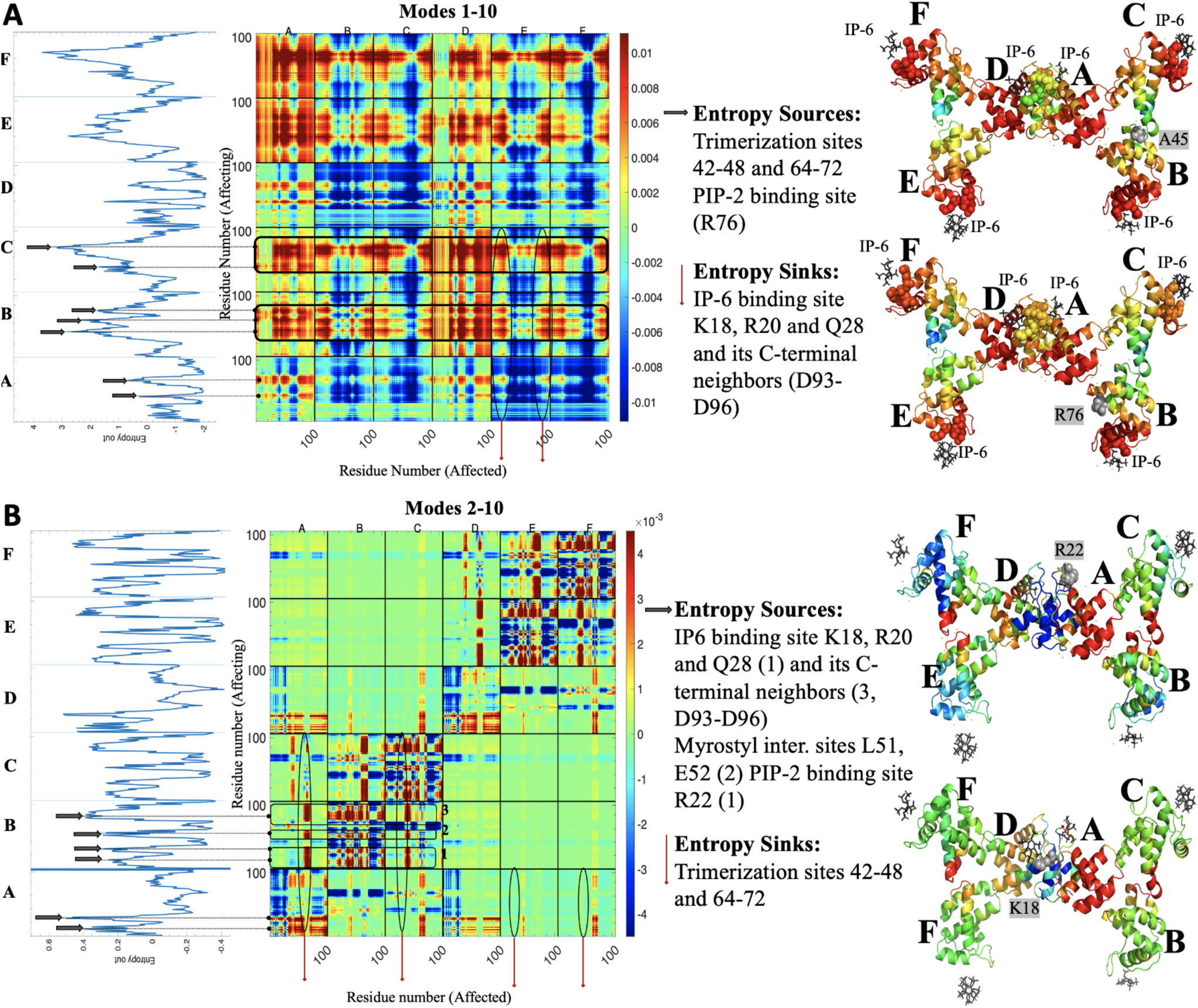
Net transfer entropy (nTE) map of hexamer (MA_IP6_C2) showing the causal interrelation for each residue pair (affecting versus affected) calculated by using an average of ten slowest modes (A) and average two to ten slowest modes (B). Cumulative transfer entropy plots (cTEs) corresponding to each nTE map are given on the left. On the right, the causal interrelation of exemplary residues from respective nTE maps, A45 and R76 on chain B **(A)** and K18 and R22 on chain A **(B)** represented with gray spheres, with the rest of residues on hexamer is color-coded on structure from highest positive (red) to lowest negative (blue) nTE values of the relevant residue. Entropy source (on chains A, B, C **(A)** and chains A, B **(B)**) and entropy sinks (on chain E **(A)** and chains A, C, E, F **(B)**) are listed and indicated, respectively with black and red arrows.

**Supplementary Fig. 9:**
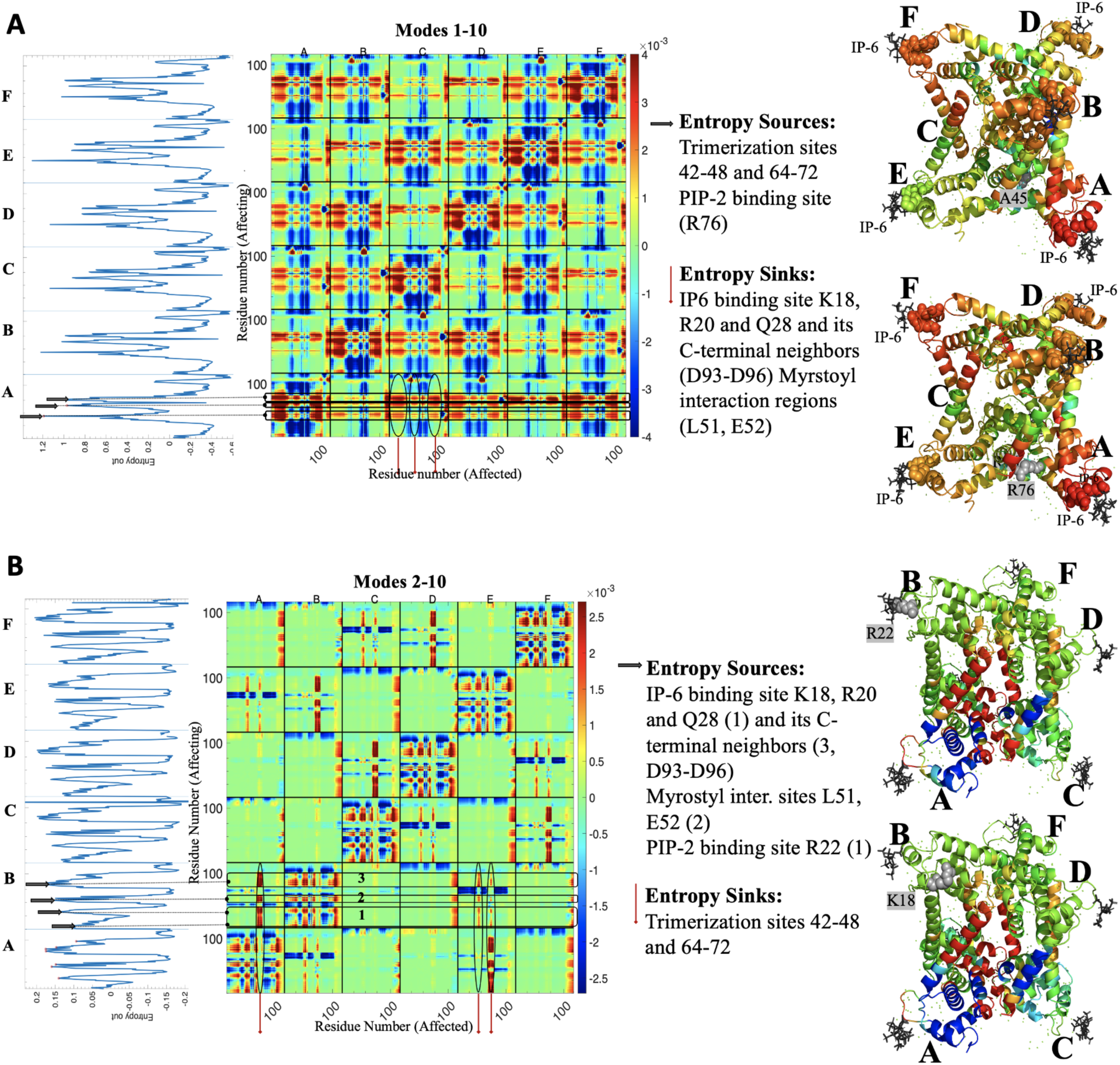
Net transfer entropy (nTE) map of hexamer (MA_IP6_R32) showing the causal interrelation for each residue pair (affecting versus affected) calculated by using an average of ten slowest modes (A) and average two to ten slowest modes (B). Cumulative transfer entropy plots (cTEs) corresponding to each nTE map are given on the left. On the right, the causal interrelation of exemplary residues from respective nTE maps, A45 and R76 on chain A **(A)** and K18 and R22 on chain B **(B)** represented with gray spheres, with the rest of residues on hexamer is color-coded on structure from highest positive (red) to lowest negative (blue) nTE values of the relevant residue. Entropy sources for chain A **(A)** and chain B (numbered 1 to 3 for IP-6 and PIP2 binding sites, myristoyl interaction sites and C-terminal IP-6 neighbors respectively) **(B)** and entropy sinks for chain C **(A)** and chains A and E **(B)** are listed and indicated respectively with black and red arrows.

**Supplementary Fig. 10:**
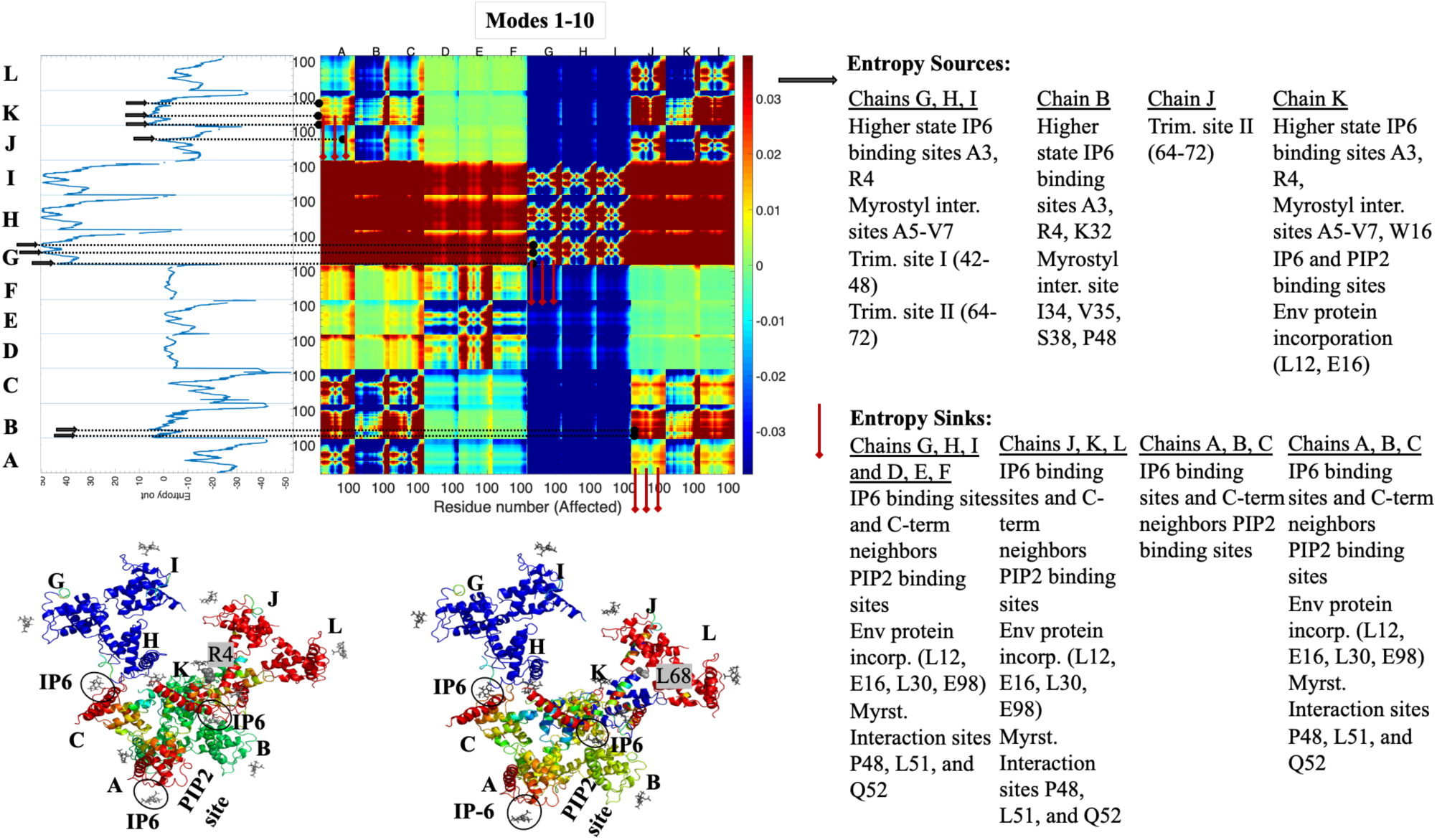
Net transfer entropy (nTE) map of dodecamer (MA_IP6_P1) showing the causal interrelation for each residue pair (affecting versus affected) calculated by using an average of ten slowest modes. Cumulative transfer entropy plots (cTEs) corresponding to each nTE map are given on the left. Below the nTE map, as exemplary cases, the causal interrelations of R4 and L68 (chain K and J respectively, gray spheres) with the rest of the residues on dodecamer are color-coded from the highest positive (red) to lowest negative (blue) nTE values. Entropy sources on chains B, G, J and K and entropy sinks on chain J are listed and indicated respectively with black and red arrows.

**Supplementary Fig. 11:**
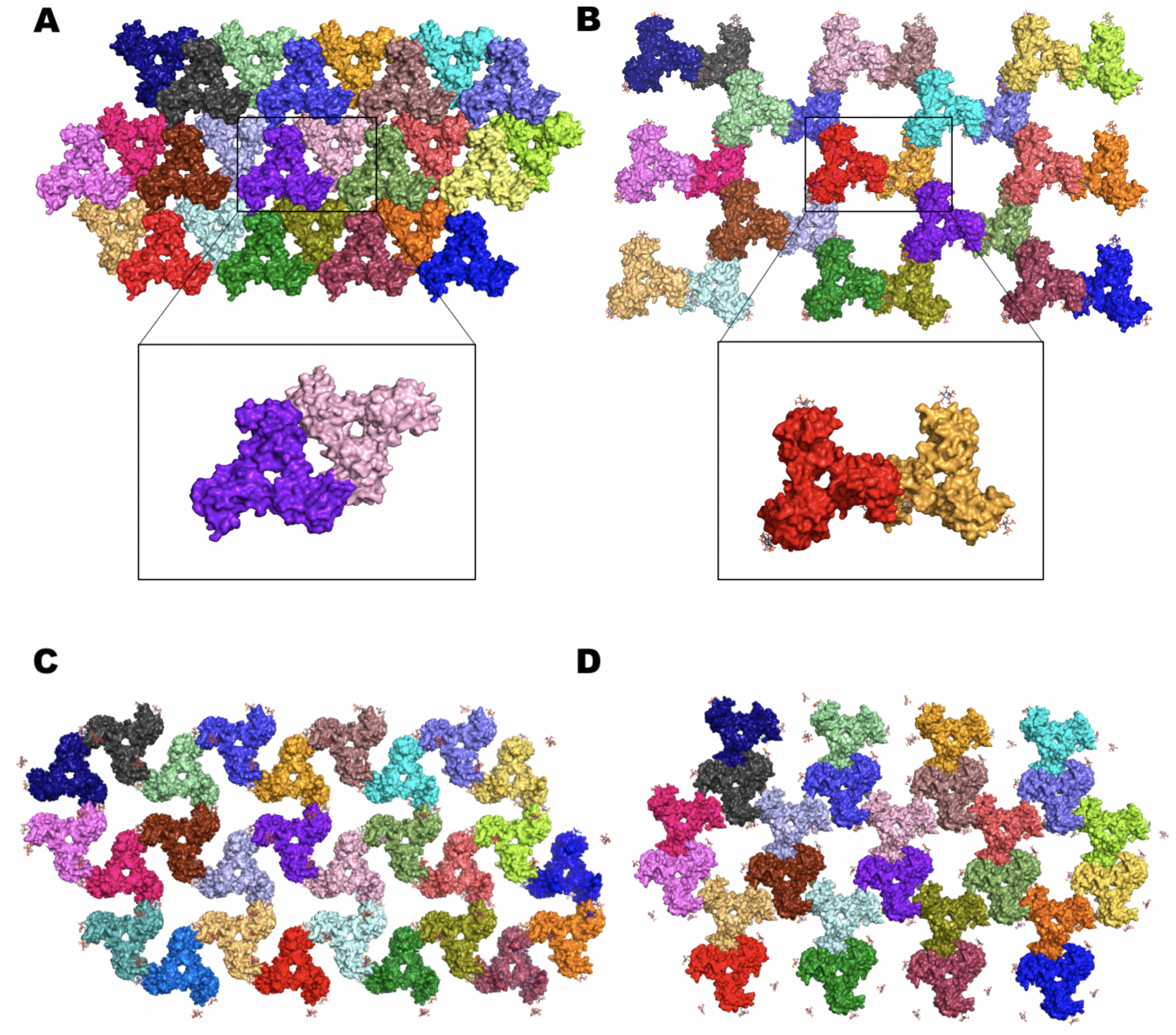
Comparison of positions of trimers in three HIV-MA structures. Three different positions of trimer are shown in 1HIW, MA_IP6_C2, MA_IP6_R32 and MA_SFX structures, respectively. **a** Estimated assembly of 1HIW structure without IP6. The asymmetric unit is shown with the purple-blue and light pink with surface representation. **b** Assembly of MA_IP6_C2 in the presence of IP6. The asymmetric units are shown with red and bright orange surfaces, respectively. **c** Trimer assembly of MA_IP6_R32 structure **d** Trimer assembly of MA_SFX structure.

**Supplementary Fig. 12:**
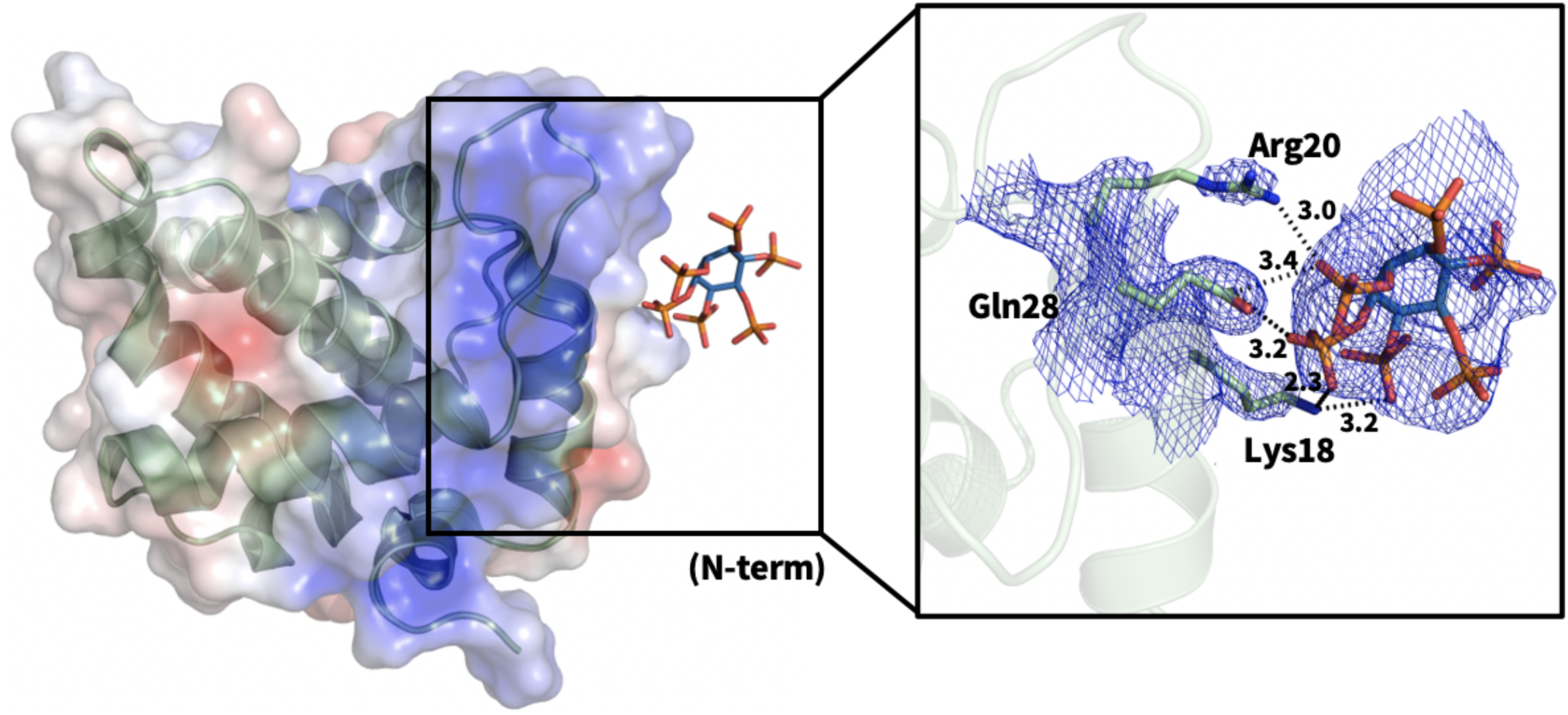
HBR within N-terminus of MA and interaction with IP6. Chain A of MA_IP6_C2 structure is colored in pale-green and carbon, oxygen and phosphorus atoms of IP6 are colored by sky-blue, red and orange, respectively with their electron density map. Polar contacts are shown with dotted lines in Angstrom. The surface of the MA_IP6_C2 structure is represented by using APBS electrostatic. Basic regions are represented by blue, acidic regions by red and hydrophobic surfaces are colored in gray.

**Supplementary Fig. 13:**
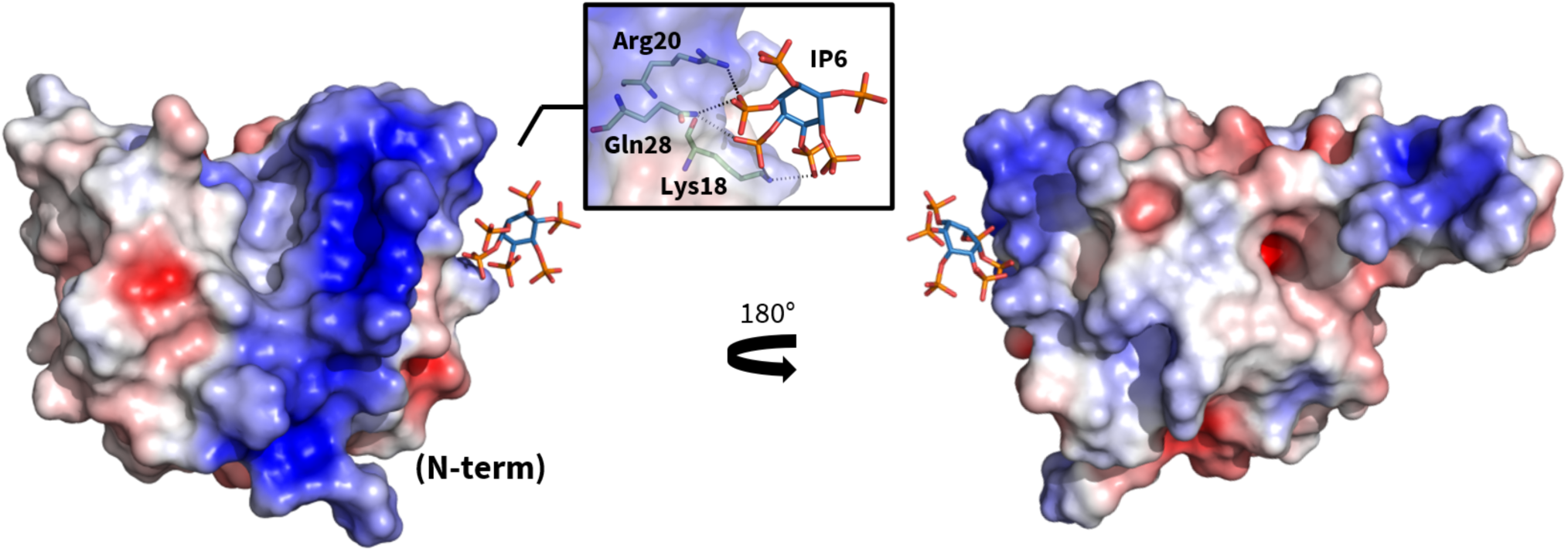
Representation of electrostatic potential surface for chain A of MA_IP6_C2 structure. APBS electrostatic is used to detect the highly basic region within the N-terminal via *PyMOL* and the structure is rotated 180 degrees in the y-axis.

**Supplementary Fig. 14:**
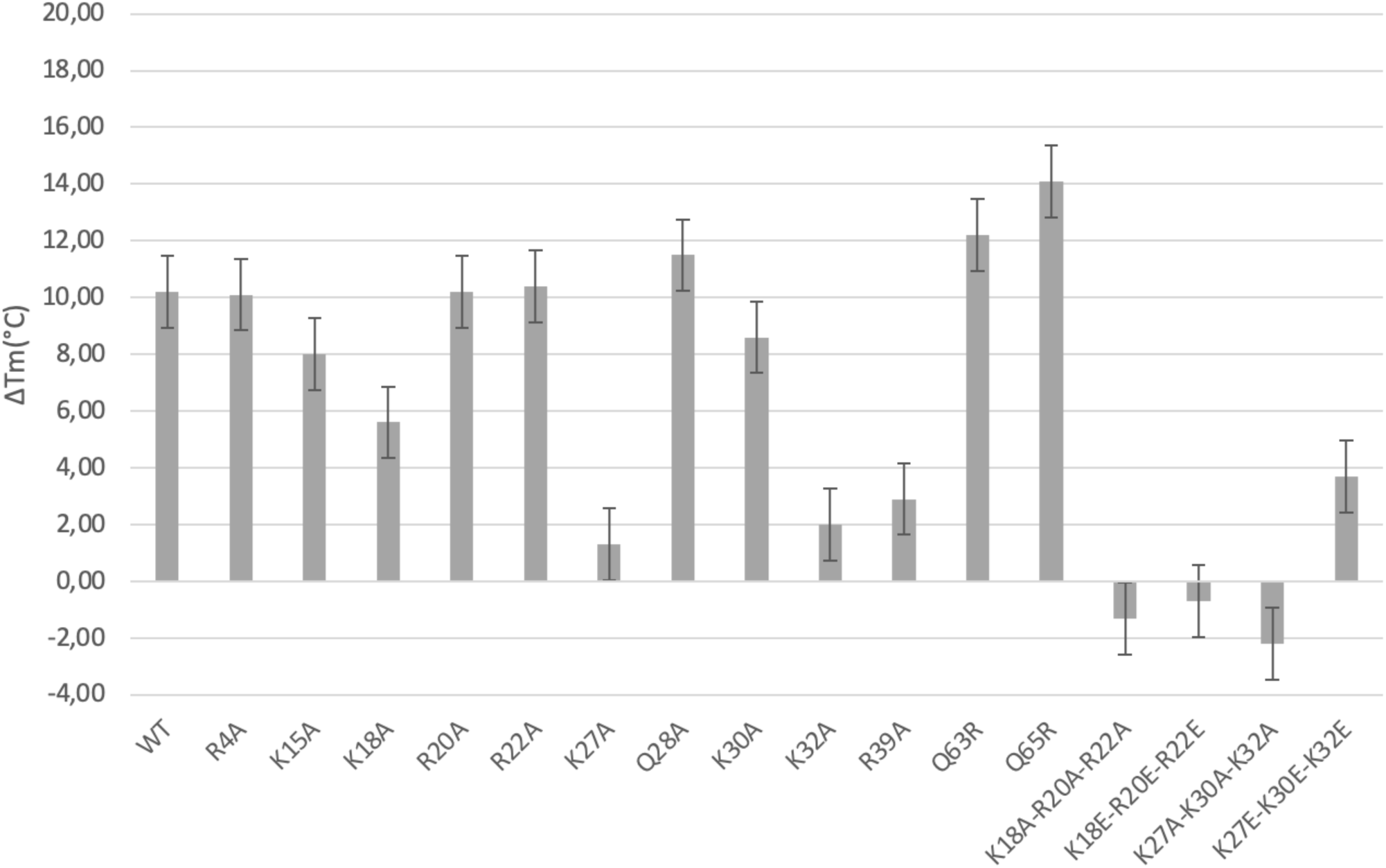
DSF assay results of point mutations of MA. The change of melting temperature (ΔTm) in the presence and absence of IP6, respectively for the wild-type and mutant MA proteins are indicated with grey bars.

## Supplementary Tables

**Supplementary Table 1:** Crystallographic Summary of MA_IP6 crystal structures.

|  | MA_IP6_R32 | MA_IP6_C2 | MA_IP6_SFX |
| --- | --- | --- | --- |
| Data Collection |  |  |  |
| Space group | *R*32 | *C*2 | *P*1 |
| Cell Dimensions |  |  |  |
| *a*, *b*, *c* (Å) | 97.83, 97.83, 176.30 | 176.18, 67.41, 97.50 | 96.60, 96.60, 91.10 |
| $\alpha$, $\beta$, $\gamma$ (°) | 90.00, 90.00, 120.00 | 90.00, 123.15, 90.00 | 90.00, 90.00, 120.00 |
| Resolution* (Å) | 39.10 - 2.40<br>(2.64 - 2.40) | 44.41 - 2.72<br>(2.82 - 2.72) | 24.57 - 3.30<br>(3.45 - 3.30) |
| *Cryo* *R*<sub>merge</sub> | 0.214 (1.54) | 0.0988 (0.799) |  |
| *XFEL* *R*<sub>split</sub> |  |  | 0.315 (6.139) |
| *I*/σ*I* | 13.21 (2.20) | 15.74 (2.42) | |
| *XFEL* *SNR* |  |  | 2.67 (0.18) |
| Completeness | 99.55 (99.53) | 99.82 (99.88) | 100.0 (100.0) |
| Redundancy | 20.3 (21.2) | 6.8 (7.0) | 62.3 (36.1) |
| CC1/2 | 0.998 (0.858) | 0.998 (0.845) | 0.958 (0.087) |
| Refinement |  |  |  |
| Resolution (Å) | 39.10 - 2.40<br>(2.64 - 2.40) | 44.41 - 2.72<br>(2.82 - 2.72) | 24.57 - 3.30<br>(3.45 - 3.30) |
| No. of reflections | 12,931 (3,027) | 25,957 (2,732) | 35,663 (2,987) |
| *R*<sub>work</sub> | 0.220 | 0.213 | 0.351 |
| *R*<sub>free</sub> | 0.276 | 0.272 | 0.408 |
| No. of Atoms |  |  |  |
| Protein | 1,920 | 5,349 | 11,415 |
| Ligand | 126 | 324 | 504 |
| Water | 97 | 69 | 80 |
| *B* factors |  |  |  |
| Protein | 48.06 | 65.11 | 174.54 |
| Ligand | 114.18 | 140.07 | 81.75 |
| Waters | 57.26 | 54.86 | 56.99 |
| RMSD |  |  |  |
| Bond length (Å) | 0.018 | 0.022 | 0.001 |
| Bond angles (°) | 1.99 | 1.96 | 0.464 |
| PDB ID | 7E1K | 7E1J | 7E1I |

**Supplementary Table 2:** Root-mean-square deviation (RMSD) among MA structures.

| RMSD (Å) of Cα atoms of chain A<sup>†</sup> |  |  |  |  |
| --- | --- | --- | --- | --- |
|  | Coordinate error<br>(Å)* | MA_IP6_R32 | 1HIW | 2HMX |
| MA_IP6_C2 | 0.37 | 0.340 | 0.274 | 1.32 |
| MA_IP6_R32 | 0.28 |  | 0.297 | 1.38 |
| 1HIW |  |  |  | 1.34 |
| 2HMX |  |  |  |  |

**Supplementary Table 3:** Root-mean-square deviation (RMSD) among MA structures.

| RMSD (Å) of Cα atoms of chain A<sup>†</sup> |  |  |  |  |
| --- | --- | --- | --- | --- |
|  | 2H3F | 2H3I | 2H3Q | 2H3Z |
| MA_IP6_C2 | 0.864 | 0.737 | 0.742 | 0.818 |

**Supplementary Table 4:** Average *B*-factor and contact area with MA of IP6 molecules.

| Objects | *B*-factor (Å<sup>2</sup>) | Contact area (Å<sup>2</sup>) |
| --- | --- | --- |
| MA | 65.08 |  |
| IP6_1 | 182.97 | 214.2 |
| IP6_2 | 120.05 | 161.1 |
| IP6_3 | 144.01 | 136.2 |
| IP6_4 | 161.55 | 155.9 |
| IP6_5 | 129.75 |  |
| IP6_6 | 117.63 |  |

